# Nuclear-import receptors remodel the dilute phase to suppress phase transitions of RNA-binding proteins with prion-like domains

**DOI:** 10.1101/2025.11.14.688546

**Authors:** Miriam Linsenmeier, Min Kyung Shinn, Thomas R. Mumford, Vicky Liu, Lukasz J. Bugaj, Rohit V. Pappu, James Shorter

## Abstract

RNA-binding proteins (RBPs) with prion-like domains, including FUS, hnRNPA1, and hnRNPA2, assemble into functional, metastable condensates that organize ribostasis, but can also transition into self-templating fibrils implicated in neurodegenerative proteinopathies such as amyotrophic lateral sclerosis (ALS). How nuclear-import receptors (NIRs) antagonize this pathological transition has remained unresolved. Here, we establish that NIRs regulate the phase behavior of prion-like cargos by remodeling the dilute phase. Quantitative analyses across length scales reveal that Karyopherin-β2 (Kapβ2) preferentially binds cargo in the dilute phase to lower the effective concentration of free RBPs thereby elevating the saturation concentration for phase separation and suppressing mesoscale clustering. ALS-linked FUS^P525L^, which binds Kapβ2 weakly, evades this regulation to form pathogenic assemblies. Thus, NIRs harness polyphasic linkage, the thermodynamic relationship between ligand binding and phase equilibria, to reshape the landscape of prion-like RBP assembly states, establishing a paradigm for how ATP-independent chaperones regulate phase behavior to prevent disease-linked aggregation.

**GRAPHICAL ABSTRACT:** 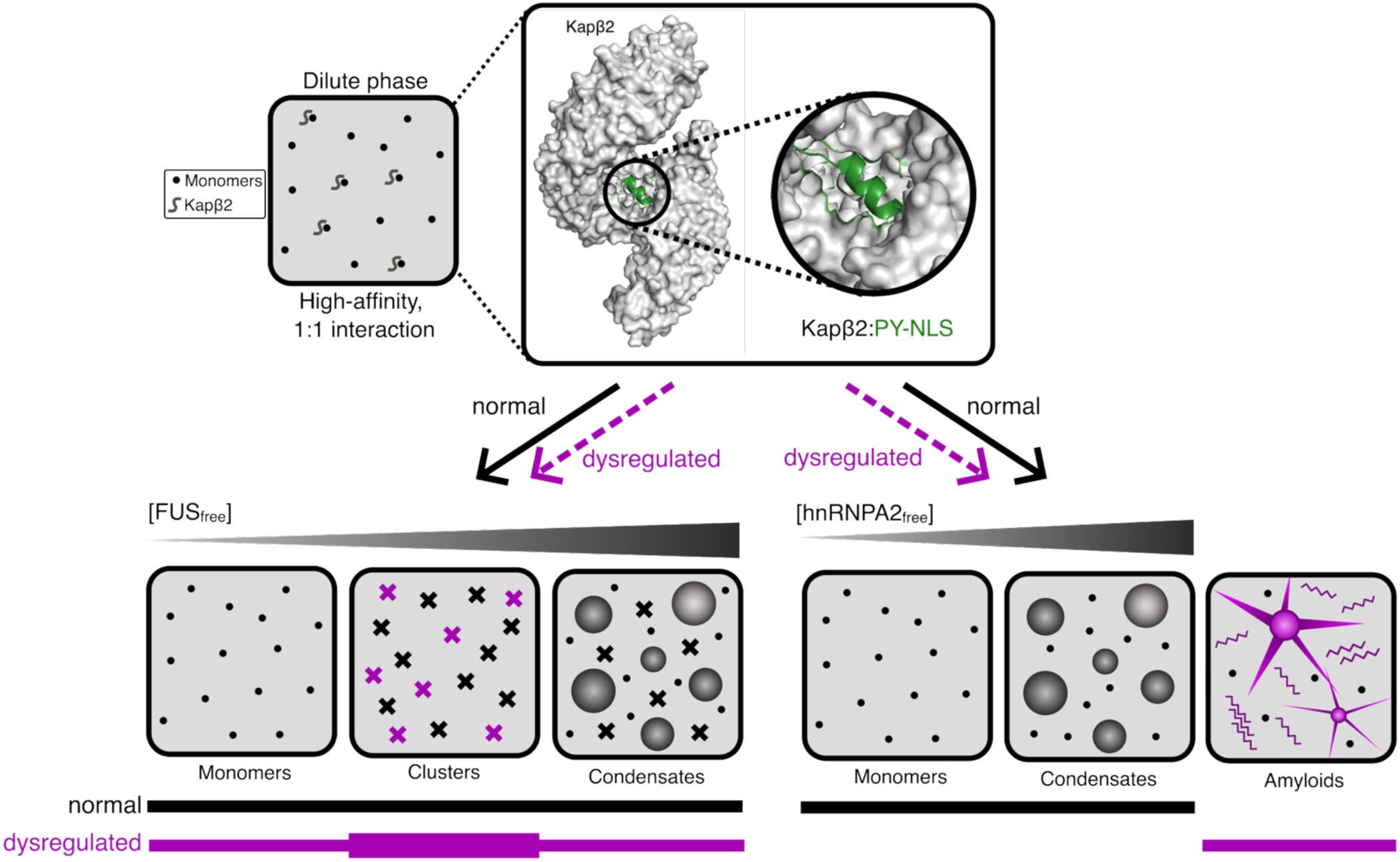

## Introduction

Cellular function relies on precise spatial and temporal control of biochemical reactions that are often organized by biomolecular condensates^1, 2^. These membraneless compartments form via phase separation of proteins and nucleic acids^1–4^. RNA-binding proteins (RBPs) with prion-like domains (PrLDs)^5–7^, such as FUS, hnRNPA1, and hnRNPA2, localize predominantly to the nucleus and are integral to the functions of various nuclear condensates^8–12^. For example, FUS is found in paraspeckles^8^, transcriptional condensates^9^, and DNA-damage-repair foci^10, 11^, whereas hnRNPA1 and hnRNPA2 assemble into 40S hnRNP assemblies^12^. Although less abundant in the cytoplasm, these RBPs also contribute to transport granules that shuttle specific mRNAs for localized translation^13–16^. Within condensates, these RBPs coordinate essential processes in RNA metabolism, including transcription, splicing, translation, transport, and degradation^2, 17^. During cellular stress, FUS, hnRNPA1, and hnRNPA2 accumulate in cytoplasmic stress granules, where they regulate mRNA triage and translation repression, and help control granule dynamics^18–21^.

Disruption of the localization and assembly of RBPs with PrLDs is tightly linked to neurodegenerative diseases^5–7, 17, 20, 22–24^. Aberrant cytoplasmic mislocalization and aggregation of FUS, hnRNPA1, and hnRNPA2 are pathological hallmarks in the brains of patients afflicted with neurodegenerative diseases such as amyotrophic lateral sclerosis (ALS) and frontotemporal dementia (FTD)^5, 20, 23, 24^. Cytoplasmic inclusions of hnRNPA1 and hnRNPA2 in degenerating muscle are also a defining feature of multisystem proteinopathy (MSP)^5, 20^. Even partial disruption of FUS nuclear localization impairs gene regulation and neuronal function, emphasizing the critical need to maintain proper localization and assembly states^25^. Moreover, recent work reveals that cytoplasmic mislocalization and aggregation of FUS in cortical neurons may contribute far more broadly to ALS cases with cognitive and behavioral impairment than previously recognized^26^.

FUS, hnRNPA1, and hnRNPA2 share a modular domain architecture that supports both self-association and RNA binding^5–7^ (**Fig. 1a**). FUS contains an N-terminal PrLD, a central RNA recognition motif (RRM), two arginine-rich (R-rich) domains flanking a zinc-finger domain, and a C-terminal proline-tyrosine nuclear localization signal (PY-NLS)^5^ (**Fig. 1a**). By contrast, hnRNPA1 and hnRNPA2 each contain two N-terminal RRMs and a C-terminal PrLD that harbors a PY-NLS^5^ (**Fig. 1a**). All three RBPs bind RNA through their RRMs, with additional contributions from arginine motifs or arginine-rich regions, and in the case of FUS the zinc-finger domain^27–31^. Multivalent interactions involving the RRMs, the PrLDs, and the R-rich regions drive phase separation via associative interactions^5, 32, 33^. Mutations in FUS, hnRNPA1, and hnRNPA2 cause neurodegenerative disease^5, 34^, and are frequently found in either the PrLD, where they alter self-association^20, 35, 36^, or the PY-NLS, where they disrupt nuclear localization^5, 23, 24, 37–42^.

**Fig. 1:**
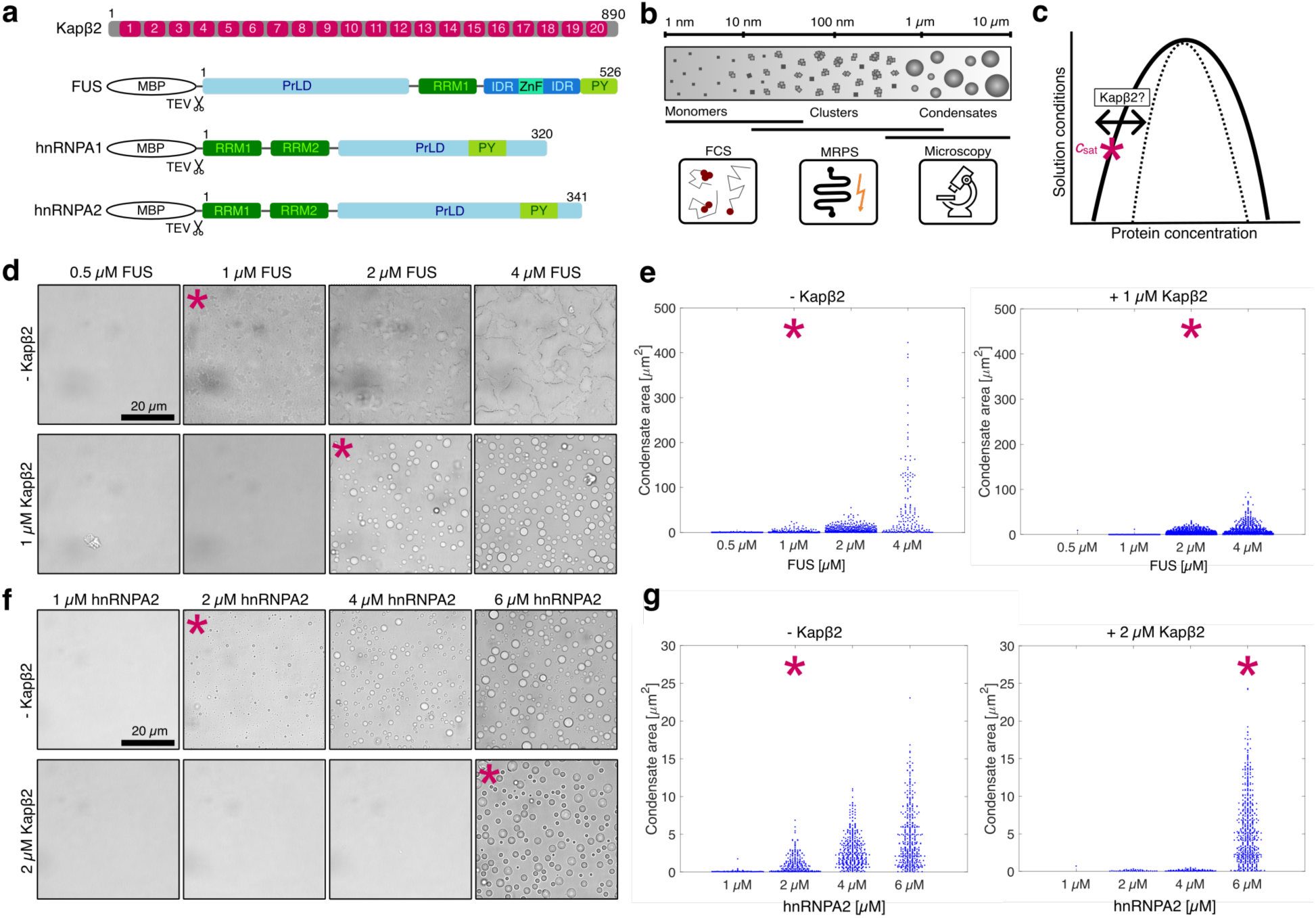
Kapβ2 increases *c*_sat_ of FUS, hnRNPA1 and hnRNPA2 due to preferential binding to cargo in the dilute phase. **a**, Domain maps of ý-importin Kapβ2 and RNA-binding proteins FUS, hnRNPA1 and hnRNPA2. Kapβ2 is composed of 20 HEAT repeats (H1-20). FUS, hnRNPA1 and hnRNPA2 are expressed with a Maltose-binding protein (MBP) tag, which is cleaved by TEV protease. Common domains are RNA-recognition motifs (RRMs), prion-like domains (PrLDs), arginine-rich intrinsically disordered regions (IDRs), and proline-tyrosine nuclear-localization sequences (PY). **b,** Techniques used to measure size distributions of higher-order protein assemblies across length scales (nm-µm range). Monomers and small clusters that are in the size range of a few nanometers to tens of nanometers were detected via Fluorescence Correlation Spectroscopy (FCS). Nano- and mesoscale clusters and small condensates that fall within the size range of 70 nm – 2 µm were probed using Microfluidic Resistive Pulse Sensing (MRPS). Larger, micron-scale condensates were imaged using widefield microscopy. **c,** Schematic phase diagram of a single-component system with the left and right arm representing the concentration of protein in the dilute and in the condensate phase, respectively. The asterisk (*) represents the saturation concentration *c*_sat_, which can be shifted by binding of a ligand to the phase separating protein. **d,** Brightfield micrographs of FUS samples at increasing concentrations in the absence (top) and presence of Kapβ2 (bottom). The intrinsic *c*_sat_ of FUS is 1 µM; in the presence of 1 µM Kapβ2, the c_sat_ of FUS increases to 2 µM. **e,** Quantification of the condensate area from images in d. **f,** Brightfield microscopy to determine the *c*_sat_ of hnRNPA2. In the absence of Kapβ2, condensates form at 2 µM hnRNPA2. In presence of 2 µM Kapβ2, the *c*_sat_ of hnRNPA2 shifts to 6 µM. **g,** Condensate area in µm^2^ extracted from images in f. Images in d and f are representative images of at least three independent replicates. Each blue dot in e and g corresponds to the area of a single condensate, and the beeswarm plot is a combination of three images. In panels d-g, the asterisks mark the *c*_sat_.

Above a saturation concentration (*c*_sat_), which depends on solution conditions, RBPs with PrLDs undergo phase separation to form micrometer-scale condensates known as macrophases^18, 32, 35, 43^ (**Fig. 1b**). The condensate interface delineates the dense and dilute phases and serves as a critical site for biochemical reactions^44, 45^. However, interfaces can promote the conversion of RBPs into pathogenic amyloid fibrils linked to neurodegenerative disease^46–49^. Indeed, synthetic FUS fibrils possess prion-like properties that seed pathology and drive neurodegeneration *in vivo*^50^. Beyond condensates and fibrils, FUS, hnRNPA1, and hnRNPA2 also form heterogeneous nanoscale and mesoscale clusters under subsaturated conditions^51–53^ (**Fig. 1b**). These clusters evolve in size and abundance with protein concentration, lowering the barrier for condensate formation while also facilitating localized RNA processing^43, 51, 52, 54^. Collectively, clusters, condensates, and fibrils define a continuum of higher-order assemblies with distinct consequences for cellular function and disease.

We propose that maintenance of the proper balance across this continuum of assemblies is essential for cellular health. Misregulation can lower the metastability of condensates formed by RBPs with PrLDs, which can lower the free energy barrier for forming stable fibrils capable of causing disease^18, 32, 35, 50, 55–58^. To counter this transition, cells deploy molecular chaperones that prevent or reverse aberrant assembly^59^. Among these chaperones are the nuclear-import receptors (NIRs), including Karyopherin-β2 (Kapβ2, also known as Transportin-1)^60–68^. Kapβ2 comprises 20 helix-turn-helix HEAT repeats (**Fig. 1a**), and binds RBPs bearing a PY-NLS with nanomolar affinity^69,^_70_. Beyond a canonical role in nuclear import^69, 71, 72^, Kapβ2 acts as a potent ATP-independent chaperone that engages cognate cargos to maintain solubility in the cytoplasm^59–64, 73–75^. Once bound, cargo is transported into the nucleus, where Ran-GTP binds Kapβ2 and triggers cargo release, completing the import cycle^60, 69, 76^.

Mutations that impair Kapβ2–RBP interactions compromise proteostasis and promote the accumulation of toxic assemblies^60–62, 70^. Such disruptions underlie early-onset or severe forms of ALS and related disorders^23, 24, 37–42^. Despite clear functional and therapeutic importance, the mechanism by which Kapβ2 modulates RBP phase behavior and suppresses aggregation remains poorly understood. Unlike canonical ATP-dependent chaperones, Kapβ2 lacks ATPase activity, raising a longstanding question: how can an ATP-independent chaperone exert precise control over the complex self-assembly landscapes of multiple prion-like RBPs?

Here, we define the mechanism by which Kapβ2 modulates the complex assembly landscapes of FUS, hnRNPA1, and hnRNPA2. Guided by the thermodynamic framework of polyphasic linkage^77–79^, we show that Kapβ2 binds prion-like RBP cargos preferentially in the dilute phase with high specificity and affinity, thereby depleting self-association-competent RBPs and shifting the phase boundary. This ATP-independent mechanism enables Kapβ2 to determine assembly outcomes irrespective of the underlying self-assembly pathway. Crucially, we directly link the NIR–RBP binding equilibrium to changes in phase equilibrium, establishing a quantitative foundation for how binding modulates condensation.

## Results

### Presence of Kapβ2 increases saturation concentrations via preferential binding to cargo in the dilute phase

To determine how Kapβ2 influences phase separation of FUS, hnRNPA1, and hnRNPA2, we measured the saturation concentration (*c*_sat_) for each RBP in the absence or presence of Kapβ2 (**Fig. 1c**). Condensate formation was initiated by TEV protease cleavage to remove the solubilizing MBP tag. Brightfield microscopy revealed that FUS condenses at ∼1 µM in the absence of Kapβ2, but this threshold increases to ∼2 µM in the presence of 1 µM Kapβ2 (**Fig. 1d, e**). hnRNPA2 shows a similar trend: *c*_sat_ increases from ∼2 µM to ∼6 µM in the presence of 2 µM Kapβ2 (**Fig. 1f, g**). For hnRNPA1, the presence of 2 µM Kapβ2 raises *c*_sat_ from ∼4 µM to ∼6 µM (**Extended Data Fig. 1a, b**). Thus, Kapβ2 consistently weakens the driving forces for phase separation of all three cargos as evidenced by the increases in threshold concentrations required for condensate formation.

Changes in the *c*_sat_ in presence of a ligand reveal how ligands interact with target macromolecules, as described by the framework of polyphasic linkage^77–79^. Equalizing the chemical potential of the RBP in the presence of a ligand L, leads to the following relationship: *c*_sat,L_ = *c*_sat,0_ (*P*_dil_/*P*_den_), where *c*_sat,L_ is the saturation concentration in the presence of the ligand and *c*_sat,0_ is the saturation concentration in the absence of the ligand. *P*_den_ is the binding polynomial that describes site-specific binding of the ligand to the macromolecules in the dense phase, and *P*_dil_ is the binding polynomial that describes site-specific binding of the ligand to the macromolecules in the dilute phase. If *P*_dil_ > *P*_den_, then binding to macromolecules in the dilute phase is preferred, and *c*_sat,L_ > *c*_sat,0_. The converse is true when ligand binding to macromolecules in the dense phase is preferred. If *P*_dil_ = *P*_den_, then binding to the macromolecules in both phases is equivalent, and *c*_sat,L_ = *c*_sat,0_. Thus, preferential binding can be inferred from how *c*_sat_ shifts upon ligand addition. Kapý2 interacts with cargo proteins through a site-specific, high-affinity interaction via the PY-NLS^61, 70, 80, 81^. We find that Kapβ2 binding consistently increases *c*_sat_, albeit to different extents for FUS, hnRNPA1, and hnRNPA2 (**Fig. 1**, **Extended Data Fig. 1**), indicating that Kapβ2 binds preferentially to these RBPs in the dilute phase to weaken the driving forces for phase separation.

Preferential binding to the dilute phase does not imply exclusion of the ligand from the dense phase. Indeed, Alexa647-labeled Kapβ2 partitions readily into the interiors of FUS and hnRNPA2 condensates (**Extended Data Fig. 2**). This observation aligns with previous reports showing that condensate partitioning and site-specific preferential binding effects are not necessarily correlated^77^. Partition coefficients reflect the contributions of multiple factors, including the physicochemical properties of the condensate environment, and therefore they do not reliably predict how ligands affect phase behaviors^77, 82^. These findings underscore the importance of the polyphasic linkage framework and the importance of assessing how *c*_sat_ is affected by ligands. Such measurements provide a direct assessment of the thermodynamic linkage between ligand binding and phase equilibria of condensate-forming systems.

### Kapβ2 remodels the size distributions of FUS assemblies that form across length scales

To assess how Kapβ2 influences higher-order FUS assemblies, we mapped the complete FUS self-assembly landscape. We quantified the size distributions of all assemblies formed in subsaturated solutions (*c* < *c*_sat_) using microfluidic resistive pulse sensing (MRPS). In MRPS, samples pass through a microfluidic cartridge containing a nano constriction under an applied electric field; each particle crossing the constriction generates a transient voltage drop proportional to particle volume^83–85^. By using devices with different constriction widths, MRPS directly measures assemblies spanning nanometer to micrometer scales, providing robust, quantitative, and unbiased size distributions for polydisperse samples. This approach thus enables comprehensive analysis of how Kapβ2 remodels FUS organization across the entire spectrum of length scales.

We first measured FUS size distributions across a range of concentrations. With brightfield microscopy we were unable to detect any visible assemblies below the *c*_sat_ of ∼1 µM (**Fig. 2a**). In contrast, MRPS revealed the presence of abundant nano- and mesoscale clusters in subsaturated solutions. The sizes of clusters, which vary depending on the degree of subsaturation (defined by the gap between *c* and *c*_sat_) were found to span tens to hundreds of nanometers in effective diameter (**Fig. 2b, Suppl. Fig. 1**). The resulting size distributions were heavy-tailed (**Fig. 2b**). These findings indicate that FUS has a strong intrinsic propensity to self-associate, favoring higher-order organization over the monomeric state even under subsaturated conditions.

**Fig. 2:**
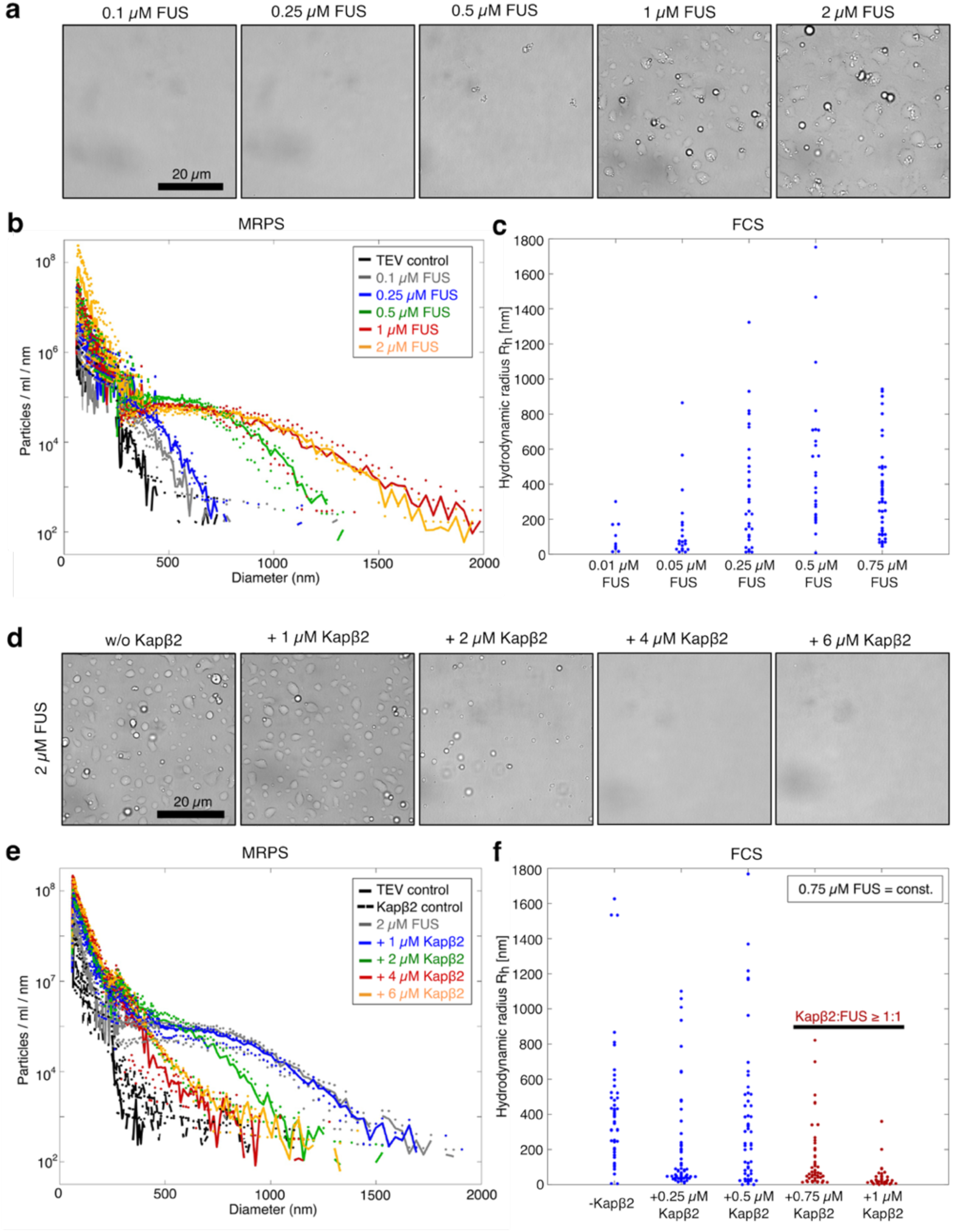
Kapβ2 remodels the totality of higher-order assemblies formed by FUS across length scales. **a**, Brightfield microscopy of samples at increasing concentrations of FUS. Condensates form above 1 µM. **b,** Size distributions of samples at increasing FUS concentrations measured via Microfluidic Resistive Pulse Sensing (MRPS). Size distributions exhibit heavy-tailed character. **c,** Particle sizes at subsaturated FUS concentrations (100 % FAM-labeled) measured by Fluorescence Correlation Spectroscopy (FCS). With increasing concentrations, larger assemblies form, covering an increasing range of sizes. **d,** Brightfield micrographs of samples at constant FUS concentrations of 2 µM in absence and presence of increasing Kapβ2 concentrations. Condensate formation is inhibited at equimolar and higher ratios of Kapβ2 to FUS. **e,** MRPS size distributions of samples at 2 µM FUS and increasing concentrations of Kapβ2. Increasing amounts of Kapβ2 shift particle sizes to smaller diameters. **f,** Particle sizes measured via FCS at 0.75 µM FUS (15 % FAM-labeled FUS) in the absence and presence of increasing concentrations of Kapβ2. With increasing concentrations of Kapβ2, small assemblies (< 100 nm, highlighted in red) are stabilized. Micrographs in a and d are representative images. Size distributions in b and e depict the mean particle counts / ml / nm of three independent replicates; dots represent the raw spread of the replicates.

Size distributions are governed by the probability *p*(*d*) of forming assemblies of a given diameter (*d*). The free energy of transitioning into a cluster of diameter *d* will be proportional to log(*p*(*d*)), and therefore the probability distribution is a direct readout of the free-energy landscape of protein self-assembly. Closer inspection of the FUS size distributions at increasing concentrations uncovered an atypical intermediate regime where the distributions flatten. Assemblies in this range (∼400–1100 nm) occur with comparable probability, consistent with a local flattening of the energy landscape^86, 87^. As a result, FUS populates a polydisperse ensemble of metastable, mesoscale clusters. Notably, a similar flattening trend has been observed in cells for the self-assembly of the nuclear elongation factor NELF^88^. This behavior deviates from continuously decaying distributions characteristic of classical aggregation driven by nucleation and growth^89^.

MRPS cannot detect assemblies that have effective diameters smaller than ∼70 nm. Therefore, we turned to fluorescence correlation spectroscopy (FCS) to probe FAM-labeled FUS at subsaturated concentrations. In the absence of TEV protease, MBP–FUS remained below 100 nm across all concentrations (**Suppl. Fig. 2**). Upon MBP removal by TEV cleavage, a continuum of assemblies emerged, with sizes increasing progressively with FUS concentration (**Fig. 2c and Suppl. Fig. 3**). These results demonstrate that well below *c*_sat_, FUS forms polydisperse mixtures of nano- and mesoscale assemblies that grow continuously with increasing concentration. Across the entire concentration range tested, FUS populates a continuum of assembly states ranging from monomers ∼ 4 nm) to nano- and mesoscale clusters (tens to hundreds of nanometers) and condensates (>1 µm) above *c*_sat_ (**Fig. 2a–c**). The heavy-tailed size distributions indicate that small assemblies predominate, while progressively larger assemblies become increasingly rare.

Next, we asked whether Kapβ2 modulates the entire spectrum of higher-order FUS assemblies. Thus, we fixed the total FUS concentration (FUS_tot_) at 2 µM (which is above *c*_sat_), and measured size distributions across increasing Kapβ2 concentrations using brightfield microscopy, MRPS, and FCS. At equimolar concentrations of Kapβ2, condensate formation was almost completely suppressed (**Fig. 2d, e and Suppl. Fig. 4**). As Kapβ2 concentration increased further, assemblies progressively shifted toward smaller sizes, and the distributions lost their heavy-tailed character (**Fig. 2e**), resembling those observed at lower FUS concentrations. At high Kapβ2 concentrations, the distributions exhibited a strictly decaying form, indicating that Kapβ2 determines whether the intermediate regime corresponding to mesoscale clusters can emerge (**Fig. 2e**). These findings suggest that Kapβ2 remodels the free energy landscape of FUS self-assembly, stabilizing the dilute phase to prevent the transition toward macroscopic condensation.

To identify the regime in which Kapβ2 most effectively modulates FUS self-assembly, we quantified the Kullback–Leibler (KL) divergence^90, 91^ between MRPS-derived cluster-size distributions for 2 µM FUS alone and those obtained at increasing Kapβ2 concentrations (**Suppl. Fig. 5, see equation 4 in Methods**). Higher KL divergence values indicate greater differences between the compared distributions. The KL divergence rose sharply at low Kapβ2 concentrations and reached a plateau at ≥ 4 µM Kapβ2, pinpointing the most effective range of Kapβ2-mediated regulation to be approximately equimolar concentrations.

To test whether Kapβ2 also modulates FUS assemblies below the MRPS detection limit (<70 nm), we used FCS to measure FAM-labeled FUS at 0.75 µM (below *c*_sat_) in the absence or presence of increasing Kapβ2 concentrations (**Fig. 2f and Suppl. Fig. 6**). As Kapβ2 concentration increased, assembly sizes progressively decreased, indicating continuous stabilization of species smaller than 100 nm (**Fig. 2f**). At higher Kapβ2:FUS ratios (≥1:1), particles below 10 nm became detectable, likely corresponding to monomeric FUS and FUS–Kapβ2 complexes. These results demonstrate that Kapβ2 preferentially stabilizes smaller species of FUS, which establishes a thermodynamic linkage whereby the stabilization of smaller species causes a destabilization of larger species via linked equilibria.

### Kapβ2 regulates cytoplasmic FUS condensation and clustering in cells

Building on mechanistic reconstitutions of the FUS–Kapβ2 system, we next examined whether FUS self-assembles through a continuum of clusters and condensates in cells and whether Kapβ2 regulates these assemblies (**Fig. 3a**). Kapβ2 engages and chaperones cognate cargos in the cytoplasm, but these interactions are disrupted in the nucleus when Ran-GTP binds and induces cargo release^60, 61^. To investigate the chaperone activity of Kapβ2 within the cytoplasm, we fused a nuclear export sequence (NES) to the C-terminal end of GFP–FUS, promoting accumulation of GFP–FUS–NES in the cytoplasm of HEK293T cells. This design preserves the interaction between FUS and Kapβ2 because the PY-NLS remains unaltered. Live-cell fluorescence microscopy and quantification of nucleocytoplasmic ratios confirmed predominant cytoplasmic localization of GFP–FUS–NES (**Fig. 3b, c**).

**Fig. 3:**
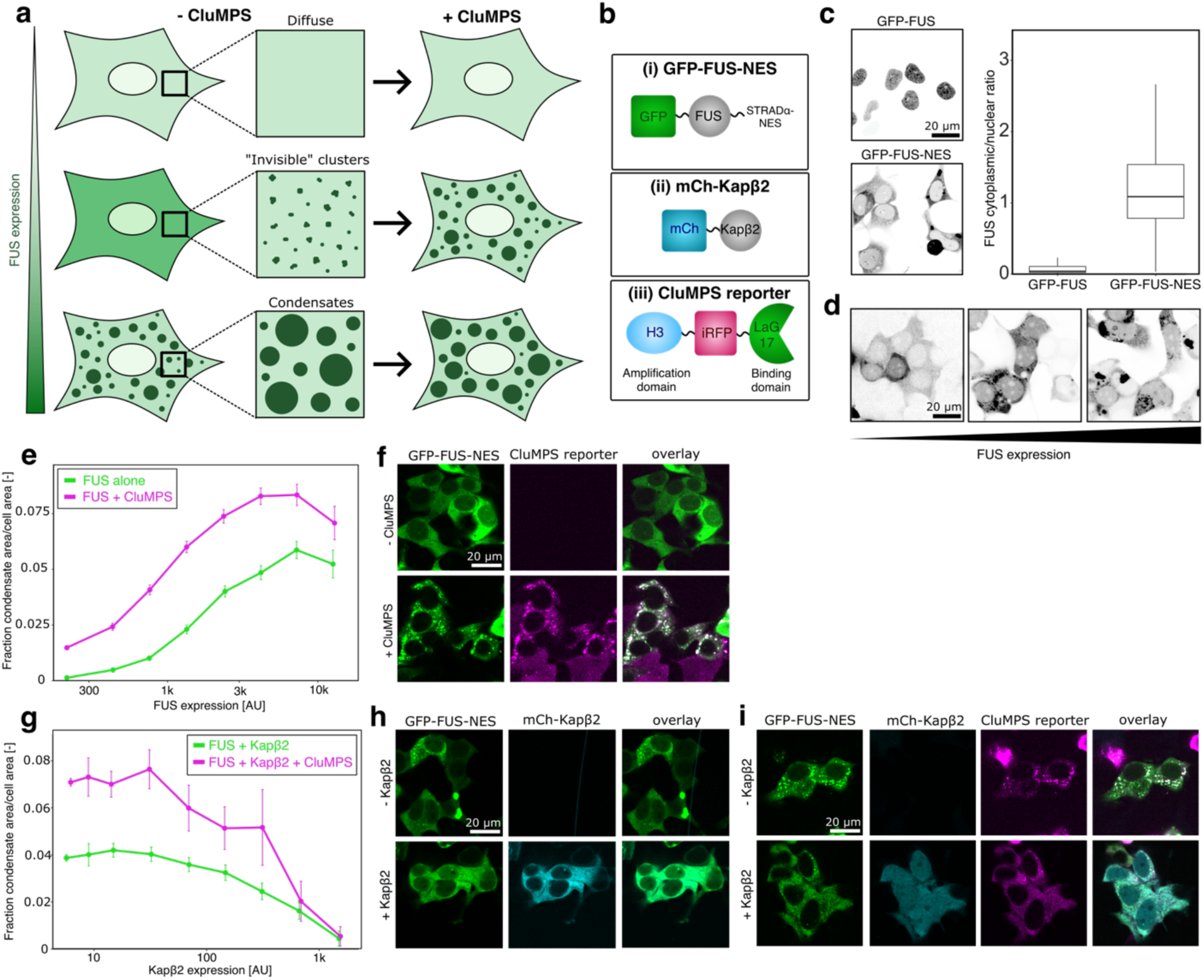
Kapβ2 modulates FUS self-assembly across length scales in cells. **a**, CluMPS technique to visualize cytoplasmic, nano- and mesoscale FUS clusters in HEK293T cells. Interaction of the CluMPS reporter with sub-diffractive FUS clusters induces the formation of CluMPS / FUS co-condensates detectable by conventional microscopy. Only cells with small assemblies exhibit diffuse FUS distribution in presence of the CluMPS reporter. This approach allows to map concentration-dependent clustering and condensation of FUS inside cells. **b,** Constructs used in this study: (i) FUS was fused to a GFP fluorophore at the N-terminus and a STRADα nuclear-export sequence (NES, sequence: GIFGLVTNLEELEVD) to the FUS C-terminal end which drives FUS to the cytoplasm. (ii) Fluorescent protein mCherry was fused to the N-terminal end of Kapβ2. (iii) The CluMPS (Clusters Magnified by Phase Separation) reporter consists of an anti-GFP nanobody (LaG17, binding domain), iRFP, and a HOTag3 oligomerization domain (H3). **c,** Confocal fluorescence microscopy and quantification of the ratio of cytoplasmic / nuclear FUS demonstrates cytoplasmic localization of the GFP-FUS-NES construct, and primarily nuclear localization of GFP-FUS. **d,** Representative fluorescence microscopy images of cells expressing increasing amounts of FUS. Higher FUS expression levels result in increased cytoplasmic condensation. **e,** Quantification of condensate area / size of cell. FUS alone does not form condensates at low expression levels but condenses at higher expression levels. In presence of CluMPS, FUS condensates can be detected at lower expression levels, reporting on nano- and mesoscale clusters forming at sub-saturated FUS expression levels. **f,** Representative fluorescence micrographs of HEK293T cells expressing FUS in absence and presence of the CluMPS reporter. **g,** Co-expression of Kapβ2 with FUS inhibits cytoplasmic FUS condensation in the absence and presence of CluMPS in a concentration-dependent manner. **h,** Representative micrographs of GFP-FUS alone and in presence of Kapβ2, demonstrating the inhibitory effect of Kapβ2 on FUS condensation. **i,** Representative images of cells co-expressing FUS with the CluMPS reporter showing that Kapβ2 inhibits FUS cluster and condensate formation in cells.

Condensates can be visualized by confocal microscopy in live cells, but nano- and mesoscale clusters fall below the diffraction limit and remain undetectable by this method. To overcome this limitation, we used Clusters Magnified via Phase Separation (CluMPS), a strategy that enlarges pre-existing clusters to enable visualization with standard fluorescence microscopy^92^ (**Fig. 3a, b**). We designed a CluMPS reporter consisting of a nanobody against GFP (LaG17, binding domain) fused to iRFP and a HOTag3 oligomerization domain (**Fig. 3b**). When co-expressed, this construct binds specifically to GFP–FUS through the nanobody–GFP interaction. At expression levels where FUS remains monomeric or forms small assemblies (up to approximately three or four FUS monomers), fluorescence from both FUS and the CluMPS reporter is expected to be diffuse throughout the cell. By contrast, when FUS forms sub-diffractive nano- or mesoscale clusters, the CluMPS reporter engages these assemblies, increases valency, and triggers measurable phase separation of both the reporter and FUS (**Fig. 3a**). To assess FUS assembly across a range of intracellular concentrations, we leveraged variable transfection efficiency, which generates a gradient of expression levels across the cell population.

Through this approach, we can directly correlate the FUS expression level with FUS assembly status within a cell and infer a minimum of three regimes. (1) FUS is monomeric or forms small clusters: diffuse distribution of FUS in absence and presence of the CluMPS reporter; (2) FUS forms nano- and mesoscale clusters: diffuse distribution of FUS in absence of the CluMPS reporter but condensation in presence of CluMPS; (3) FUS forms condensates (> *c*_sat_): detection of condensates in the absence and presence of the CluMPS reporter (**Fig. 3a**).

We first examined how FUS self-assembly depends on expression level (**Fig. 3d-3f**). Without the CluMPS reporter, FUS condensation became progressively more prominent as expression increased (**Fig. 3d**). We quantified this effect by extracting the fraction of the condensate area per cell area across a large population of cells, spanning several orders of magnitude of FUS expression levels (**Fig. 3e**, **3f, Suppl. Fig. 7a**). At low expression, FUS appeared diffuse throughout the cytoplasm (**Fig. 3e, 3f**). However, in cells expressing comparable levels of FUS together with the CluMPS reporter, condensates were readily detected, indicating that CluMPS uncovers the presence of nano- and mesoscale FUS clusters that form below the saturation threshold *c*_sat_ for visible condensates. At higher expression, FUS condensation was further enhanced in the presence of the CluMPS reporter (**Fig. 3e, 3f**), consistent with the coexistence of nano- and mesoscale clusters. In agreement with our biochemical reconstitutions (**Fig. 2a–2c**), these results demonstrate that FUS self-assembles in cells through a concentration-dependent continuum of clusters and condensates.

Next, we examined whether Kapβ2 co-expression inhibits FUS clustering and condensation in cells and if it did so in a concentration-dependent manner (**Fig. 3g-3i**). Similar to our pure protein studies (**Fig. 2d-2f**), we analyzed cells expressing comparable levels of FUS but varying amounts of Kapβ2 (**Suppl. Fig. 7b**). At low Kapβ2 expression, FUS condensates and clusters persisted both in the absence and presence of CluMPS (**Fig. 3g**). As Kapβ2 levels increased, FUS condensation and clustering progressively diminished. At the highest Kapβ2 concentrations, CluMPS no longer detected FUS clusters, indicating stabilization of monomeric FUS and small assemblies (**Fig. 3g**). These observations parallel our biochemical reconstitution studies, demonstrating that high Kapβ2 concentrations are required for full suppression of FUS self-association (**Fig. 2d–2f)**. Importantly, CluMPS reporter expression remained constant across all cells analyzed, confirming that the observed effects result from changes in Kapβ2 concentration rather than reporter abundance (**Suppl. Fig. 7c**). Together, these findings show that FUS can form a concentration-dependent continuum of higher-order assemblies in the cytoplasm, which is tightly regulated by Kapβ2. Thus, Kapβ2 serves as a key cytoplasmic chaperone that remodels the phase behavior of FUS to maintain solubility and prevent excessive condensation.

### High-affinity FUS-Kapβ2 interaction in the dilute phase regulates FUS assembly

Kapβ2 and FUS form a stable 1:1 complex through a high-affinity interaction between the concave surface of Kapβ2 and the PY-NLS of FUS^70, 74^. We hypothesized that this high-affinity interaction in the dilute phase controls FUS self-assembly by renormalizing the concentration of free, self-association–competent FUS (FUS_free_). Building on this idea, and guided by our results (**Fig. 1-3**), we developed a quantitative model to describe how the Kapβ2–FUS binding equilibrium influences the overall equilibria of assemblies and phase equilibrium of FUS across length scales. There are two possible scenarios to consider, and the model we developed tests one of these as the most parsimonious explanation of our findings. In model 1, the high affinity, 1:1 binding of Kapβ2 and FUS lowers the free pool of FUS that is available for the cascade of hierarchical associations that we observe using different techniques. In this linked equilibrium model, 1:1 binding affects the association and phase equilibria. Model 2 envisages a more complex scenario, whereby Kapβ2 binds differently to different species that form along the assembly pathway. To test this model, we would need to know the microscopic binding constants for each of the species. Prior to investigating complex scenarios such as model 2, we asked if a simple linked equilibrium model is sufficient to explain the totality of the data.

Our data indicate that when the total FUS concentration (FUS_tot_)) exceeds the saturation threshold (*c*_sat_), monomers, clusters, and condensates coexist in equilibrium (**Fig. 2a–2c, 4a**). Increasing Kapβ2 concentration (Kapβ2_tot_) promotes the formation of stable Kapβ2–FUS complexes in the dilute phase, thereby depleting FUS_free_. When FUS_free_ remains above *c*_sat_, monomers, clusters, and condensates all coexist. Conversely, when FUS_free_ falls below *c*_sat_, condensates do not form. For the clusters that persist, their overall abundance is altered by the renormalization of the concentration of free FUS by Kapβ2. This shift in assembly behavior is reflected in reduced assembly sizes and counts as Kapβ2_tot_ increases (**Fig. 2d, 2e**). Together, these results support the hypothesis that Kapβ2 regulates FUS self-assembly by modulating the pool of free FUS in the dilute phase, thereby determining whether FUS transitions from clusters to condensates.

To test the linked equilibrium model, we calculated the concentration of free, self-association-competent FUS_free_ as a function of Kapβ2_tot_ (**Fig. 4b; see equations 5–8 in Methods**). The dissociation constants (*K*_d_) were set to be 10, 50, 75, and 100 nM for the interaction of Kapβ2 with the isolated FUS PY-NLS peptide (amino acids 475-525). This range of *K*_d_ values was based on previously reported isothermal titration calorimetry (ITC) measurements^61, 70, 81^ and they were obtained under solution conditions similar to those used here. A recent study also reported a *K*_d_ of 43 nM for full-length FUS binding to full-length Kapβ2, measured by fluorescence anisotropy under different buffer conditions^80^, and this value was included for comparison. Consistent with our experimental setup, FUS_tot_ was fixed at 2 µM. The resulting calculations yielded theoretical estimates of FUS_free_ across increasing Kapβ2 concentrations (**Fig. 2d, 2e and Fig. 4b**), enabling quantitative evaluation of the proposed linked equilibrium model.

**Fig. 4:**
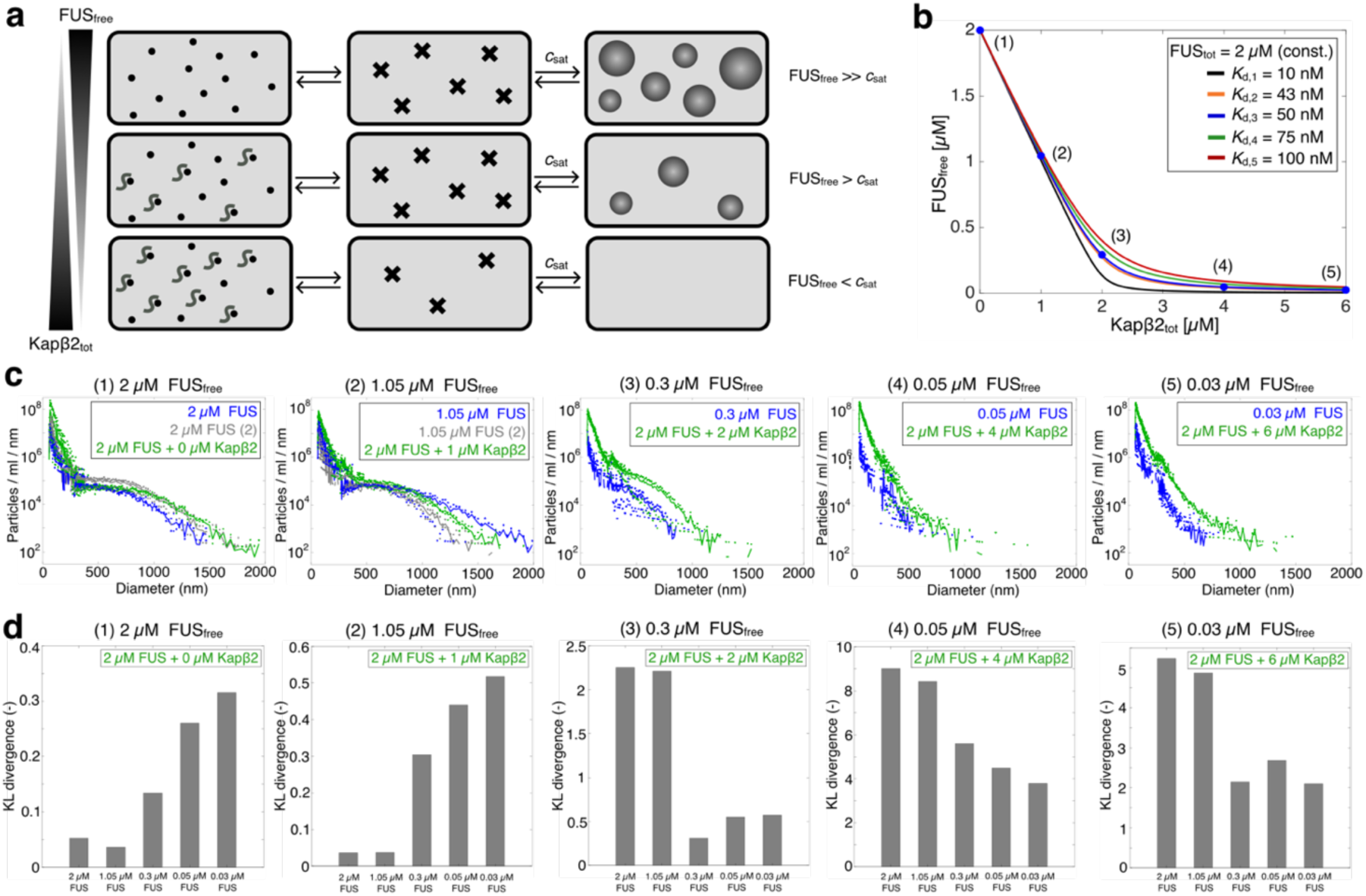
Kapβ2 regulates FUS self-assembly by controlling the concentration of free FUS (FUS_free_) in the dilute phase via a single-site, high-affinity interaction. **a**, Model of chaperone mechanism of Kapβ2. At FUS concentrations above *c*_sat_, condensates, clusters and monomers co-exist. Preferential binding of Kapβ2 FUS with high affinity in the dilute phase decreases the concentration of free, self-association-competent FUS (FUS_free_), and this suppresses the formation of higher-order structures. **b,** Theoretical estimate of the concentration of FUS_free_ as a function of the total Kapβ2 concentration (Kapβ2_tot_) derived from Equations 5-8. Conditions (1)-(5) were tested experimentally via MRPS. The total FUS concentration (FUS_tot_) was constant (2 µM) while the dissociation constant *K*_d_ of the Kapβ2:FUS complex was varied (*K*_d,1_ = 10 nM – black, *K*_d,2_ = 43 nM – orange, *K*_d,3_ = 50 nM – blue, *K*_d,4_ = 75 nM – green, *K*_d,5_ = 100 nM – red). **c,** Comparison of size distributions at 1.05, 0.3 and 0.03 µM FUS in the absence of Kapβ2, with size distributions at 2 µM FUS in presence of 1, 2 and 6 µM Kapβ2, resulting in the same concentrations of FUS_free_. Experiments correspond to conditions (1)-(5) in b, assuming a *K*_d_ of 50 nM. Size distributions in c show the mean particle count/ml/nm of three independent replicates; dots represent the raw spread of the replicates. **d,** Kullback-Leibler (KL) divergence as a measure to compare size distributions of samples at 2 µM FUS in presence of increasing concentrations of Kapβ2 with size distributions at various concentrations of FUS alone. Low KL divergence indicates high similarity between distributions with the same theoretically estimated concentration of FUS_free_.

Next, we compared size distributions from FUS–Kapβ2 mixtures with those obtained from matched concentrations of FUS_free_ alone (**Fig. 4c**). Using an intermediate *K*_d_ of 50 nM, we analyzed samples containing 2 µM total FUS with 0, 1, 2, 4, and 6 µM Kapβ2 (conditions 1–5) and compared them to FUS-only samples of 2, 1.05, 0.3, 0.05, and 0.03 µM, corresponding to the calculated FUS_free_ concentrations in the presence of Kapβ2 (**Fig. 4c**). MRPS measurements revealed strikingly similar size distributions between matched pairs, with comparable assembly sizes, counts, and heavy-tailed profiles. FCS measurements confirmed this behavior at smaller length scales (**Extended Data Fig. 3**). These results demonstrate that the linked equilibrium model accurately predicts assembly behavior across concentrations, linking thermodynamic binding parameters to the emergent size distributions of FUS assemblies.

To quantitatively assess the similarity between experimental size distributions, we calculated KL divergence values (**see Methods section**). Distributions from samples containing 2 µM total FUS and varying Kapβ2 concentrations (0, 1, 2, 4, and 6 µM) were compared with those from FUS-only samples at 2, 1.05, 0.3, 0.05, and 0.03 µM, which correspond to the calculated concentrations of FUS_free_. KL divergence values reached a minimum when the compared samples represented equivalent FUS_free_ concentrations (**Fig. 4d**), indicating maximal similarity between the two sets. This quantitative agreement confirms that the linked equilibrium model accurately predicts the redistribution of FUS between free and complexed states, capturing how binding equilibria shape the emergent self-assembly landscape.

Taken together, our analysis based on model 1 demonstrates that FUS self-assembly is fundamentally determined by the concentration of unbound, self-association–competent FUS in the dilute phase (FUS_free_). Specific, high-affinity binding of Kapβ2 to the PY-NLS of FUS generates a 1:1 complex, thereby reducing the concentration of FUS_free_. We refer to this available pool of unbound FUS as the *effective concentration* of FUS available for self-association. Lower effective concentrations weaken the concentration-dependent self-associations that drive clustering and condensate formation. Thus, preferential binding of Kapβ2 to FUS in the dilute phase modulates the effective concentration of FUS available for homotypic interactions, thereby regulating self-assembly across all length scales. Our analysis demonstrates that the effective concentration of FUS available for self-association is the key parameter determining FUS self-assembly, including the formation of a regime of mesoscale-sized networked clusters. Thus, we directly link the Kapβ2–FUS binding equilibrium established in the dilute phase to the overall FUS phase equilibrium and to the full continuum of higher-order assemblies formed by FUS.

### Impaired Kapβ2-FUS interaction results in dysregulated self-assembly of ALS-linked FUS_P525L_

Having established the sufficiency of a parsimonious biophysical mechanism by which Kapβ2 regulates self-assembly of wild-type FUS, we next asked whether this control is disrupted for the ALS-linked variant FUS^P525L^, which carries a mutation within the PY-NLS and causes aggressive, early-onset disease in patients as young as 11 years of age^42^. Substitution of proline 525 with leucine decreases the binding affinity of FUS for Kapβ2 to an estimated *K*_d_ of ∼200 nM, which is ∼4-fold weaker than that of wild-type FUS^60, 81, 93^. This reduced affinity impairs nuclear import and promotes cytoplasmic mislocalization, providing a direct link between weakened Kapβ2 engagement and pathological FUS accumulation.

We first examined how Kapβ2 influences the *c*_sat_ of FUS^P525L^ by imaging samples across increasing protein concentrations in the absence or presence of 2 µM Kapβ2 (**Fig. 5a**). Condensates formed above a *c*_sat_ of ∼1–2 µM FUS^P525L^. Similar to wild-type FUS, the addition of Kapβ2 shifted the *c*_sat_ to higher concentrations, increasing *c*_sat_ to ∼3 µM (**Fig. 5a, b**). This shift indicates that Kapβ2 preferentially interacts with FUS^P525L^ in the dilute phase, but with reduced efficacy relative to wild-type FUS, consistent with weaker binding affinity.

**Fig. 5:**
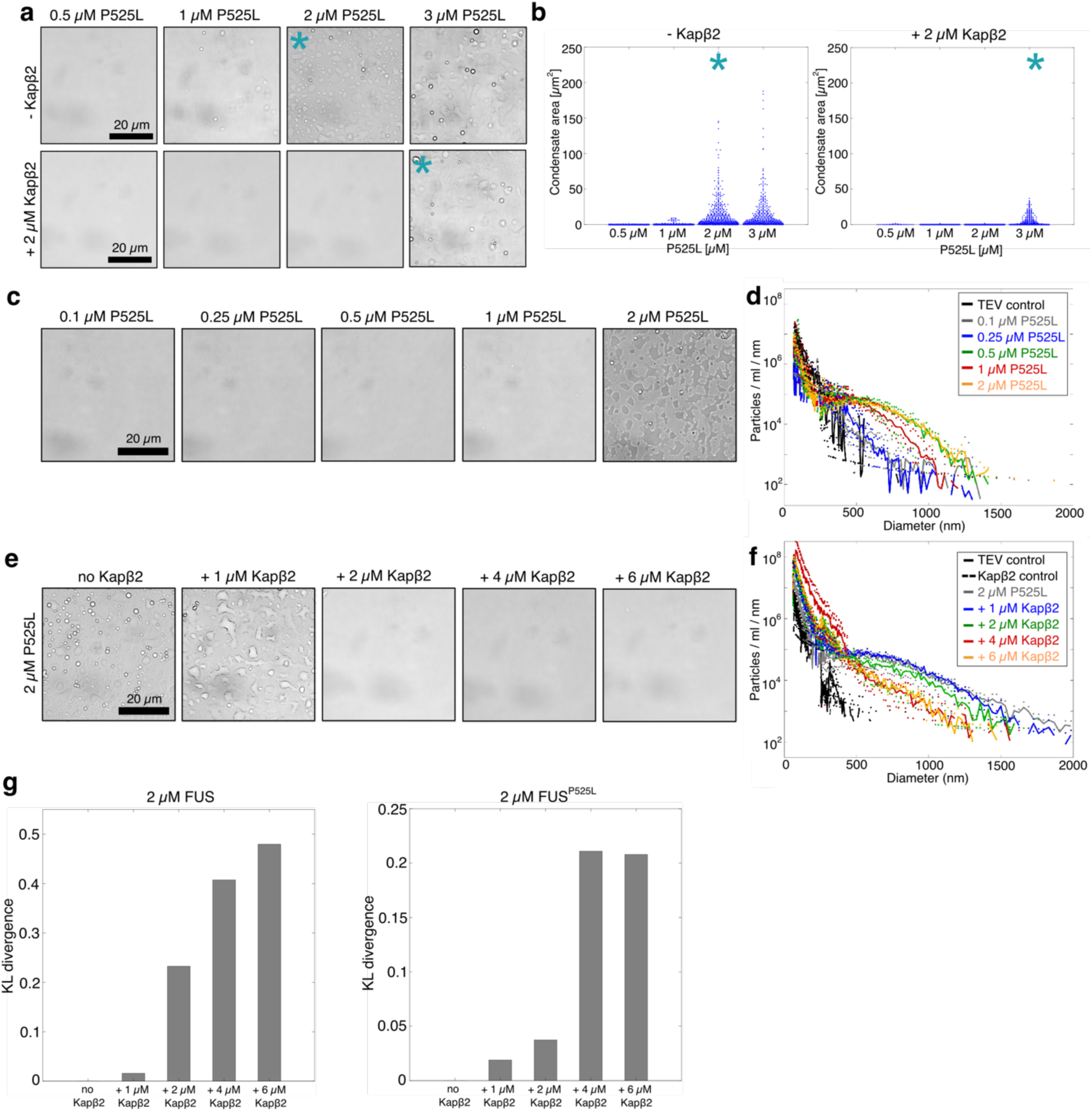
Kapβ2-mediated control of the spectrum of assembly states FUS is weakened by the ALS-associated mutation FUS^P525L^. **a**, *c*_sat_ of FUS^P525L^ in the absence and presence of 1 µM Kapβ2. The FUS^P525L^ variant exhibits a *c*_sat_ of 1-2 µM. In presence of 2 µM Kapβ2, *c*_sat_ shifts to 3 µM FUS^P525L^. **b,** Quantification of the condensate areas in µm^2^ from images in a. Blue dots represent areas of single condensates. Asterisks (*) mark the *c*_sat_. **c,** Representative brightfield microscopy images of samples formed at increasing concentrations of FUS^P525L^. Condensates form at ∼ 2 µM FUS^P525L^. **d,** Size distributions of increasing concentrations of FUS^P525L^ measured by MRPS, showing a shift to larger particle sizes with increasing concentrations. **e,** Effect of Kapβ2 on condensates formed at 2 µM FUS^P525L^. Condensate formation is inhibited at equimolar concentrations of FUS^P525L^ and Kapβ2 (2 µM respectively). **f,** Size distributions of samples at 2 µM FUS^P525L^ in the absence and presence of increasing Kapβ2 concentrations, acquired via MRPS. Kapβ2 inhibits condensate formation, but only mildly affects particle sizes and counts below *c*_sat_. Panels d and f show the mean particle counts/ml/nm of three independent replicates; dots represent the raw spread of the replicates. **g,** Kullback-Leiber (KL) divergence of distributions at 2 µM FUS or FUS^P525L^ in absence of Kapβ2 compared to distributions in presence of increasing concentrations of Kapβ2. KL divergence for FUS samples increases at lower Kapβ2 concentrations compared to FUS^P525L^ samples, indicating inefficient regulation of FUS^P525L^ self-assembly.

We next measured the size distributions of FUS^P525L^ assemblies across increasing protein concentrations using MRPS (**Fig. 5c, d and Suppl. Fig. 8**). FUS^P525L^ displayed a self-association profile broadly similar to wild-type FUS, with nano- and mesoscale clusters detected in subsaturated solutions and heavy-tailed size distributions under all conditions tested (**Fig. 5d**). Notably, FUS^P525L^ stabilized mesoscopic clusters at lower concentrations than wild-type FUS, as reflected by atypically shaped distributions appearing at 0.25 µM FUS^P525L^. These findings indicate that although FUS^P525L^ retains overall phase-separation behavior similar to wild-type FUS, the mutant displays a slightly enhanced propensity to form and stabilize nano- and mesoscale clusters well below the condensation threshold.

To evaluate how Kapβ2 modulates condensate formation and clustering of the FUS^P525L^ variant, we performed brightfield microscopy and MRPS analyses on samples containing 2 µM FUS^P525L^ (above *c*_sat_) with increasing Kapβ2 concentrations. Approximately 2 µM Kapβ2 was required to suppress condensate formation, similar to the concentration needed for wild-type FUS (**Fig. 5e**). However, Kapβ2 was markedly less effective at shifting size distributions toward smaller assemblies, as nano- and mesoscale clusters persisted at subsaturated FUS^P525L^ concentrations even at elevated Kapβ2 levels (**Fig. 5f and Suppl. Fig. 9**). These results reveal that weakened binding between Kapβ2 and FUS^P525L^ compromises the ability of Kapβ2 to enable a full remodeling of the assembly landscape, leaving residual clusters that escape regulation.

To quantitatively compare how Kapβ2 regulates wild-type FUS and FUS^P525L^, we calculated KL divergence values between size distributions of 2 µM FUS or FUS^P525L^ alone compared to distributions in the absence or presence of Kapβ2 (**Fig. 5g**). For wild-type FUS, KL divergence increased sharply with rising Kapβ2 concentrations, reflecting substantial remodeling of the size distributions. In contrast, FUS^P525L^ distributions remained similar to those observed without Kapβ2, yielding consistently low KL divergence values (**Fig. 5g**). These analyses demonstrate that reduced Kapβ2–FUS binding affinity diminishes the ability of Kapβ2 to regulate self-assembly, leading to persistent mesoscale clustering even at high chaperone concentrations. ALS-linked mutations within the PY-NLS, which weaken Kapβ2–FUS binding, thereby reduce the ability of Kapβ2 to modulate the effective concentration of FUS available for self-association and compromise regulation of self-assembly. This failure to control higher-order assembly formation drives cytoplasmic mislocalization and accumulation of clusters, which are hallmarks of FUS-linked neurodegeneration^22, 94, 95^.

For the isolated PrLD of hnRNPA1, the threshold concentration for fibril formation is lower than the saturation concentration for condensate formation, indicating that condensates are metastable with respect to fibrils^49^. Importantly, primary nucleation and fibril elongation can occur in the dilute phase^49^. Extrapolating from these findings, we propose that ALS-associated mutations that weaken Kapβ2 binding increase the effective concentration of FUS available for self-association, thereby promoting formation of self-associated species that may act as seeds to lower the free energy barrier for fibril formation in the dilute phase. Our results suggest that precise molecular-level control of FUS phase behavior is essential for maintaining cellular homeostasis, as even subtle disruptions can promote pathological aggregation.

### Kapβ2 inhibits aberrant condensate-to-amyloid transitions of hnRNPA2

Our data establish that Kapβ2 regulates the self-assembly of higher-order FUS structures both below and above the *c*_sat_. Notably, FUS condensates remain stable over several days without transitioning into amyloids^35^. We next asked whether Kapβ2 exerts a similar regulatory influence on another prion-like RBP, hnRNPA2. Both wild-type and disease-associated hnRNPA2 variants are prone to fibril formation^20, 60^. This system allowed us to test whether Kapβ2 can suppress condensate-to-amyloid transitions for other RBPs with PrLDs.

We first characterized hnRNPA2 self-assembly in the absence of Kapβ2 using brightfield microscopy and MRPS (**Fig. 6a, 6b; Suppl. Fig. 10**). Condensate formation occurred above 2 µM hnRNPA2 (**Fig. 6a**), consistent with the classical phase transition behavior. MRPS measurements revealed that below *c*_sat_ (0.5 and 1 µM), hnRNPA2 formed only small clusters and lacked the heavy-tailed size distributions observed for FUS. Heavy-tailed distributions only emerged above *c*_sat_ (**Fig. 6b**). These results indicate that under identical solution conditions, hnRNPA2 condensation proceeds through quasi-homogeneous nucleation-driven phase separation with minimal clustering in subsaturated solutions^52, 96, 97^.

**Fig. 6:**
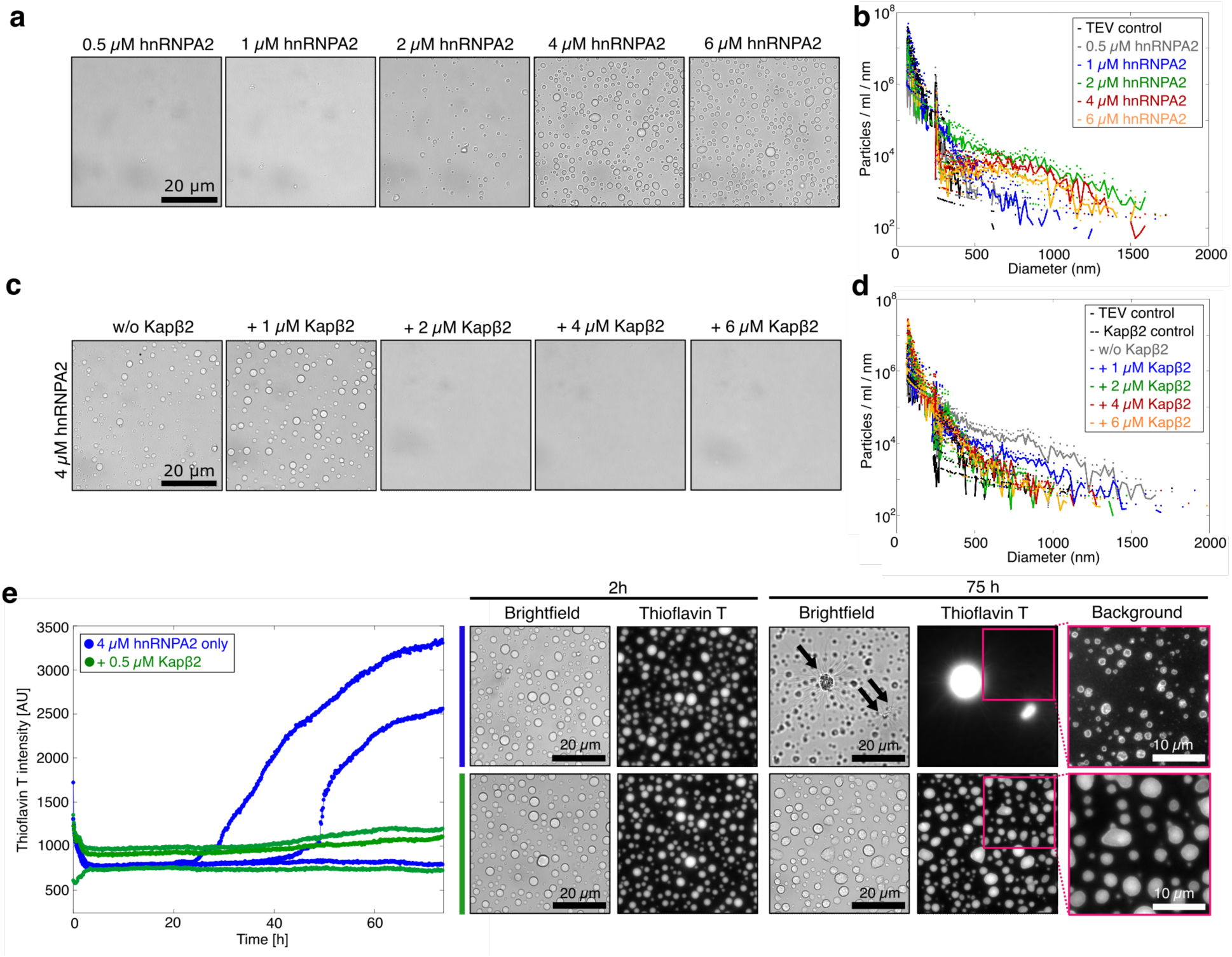
Kapβ2 regulates hnRNPA2 condensate size distributions and inhibits transition to amyloids. **a**, Representative brightfield micrographs of samples at increasing hnRNPA2 concentrations. The *c*_sat_ for condensate formation is 2 µM. **b,** MRPS-extracted size distributions of samples measured at the same conditions as in panel a. Heavy-tailed size distributions occur only above *c*_sat_ implying that the range of clusters form once the soluble phase has been saturated and not when the system is sub-saturated. The observed species likely represent those formed in the coexisting dilute phase. **c,** Brightfield microscopy of condensates at 4 µM hnRNPA2 (> *c*_sat_) in the absence and presence of increasing amounts of Kapβ2. Condensate formation is inhibited at sub-stoichiometric concentrations of Kapβ2 (2 µM). **d,** Size distributions of hnRNPA2 -/+ Kapβ2 at the same conditions as in c, measured by MRPS. Size distributions lose their heavy-tailed character at Kapβ2 concentrations which inhibit condensate formation (4 µM hnRNPA2 + 2 µM Kapβ2). Panels b and d show the mean particle counts/ml/nm of three independent replicates; dots represent the raw spread of the replicates. **e,** Thioflavin T (ThT) fluorescence signal over time of samples containing 4 µM hnRNPA2 in the absence (blue) and presence (green) of 0.5 µM Kapβ2. After 2 h of incubation, both samples form condensates, as visualized by brightfield and fluorescence microscopy and low ThT intensity. Over time, hnRNPA2 transitions to amyloids, represented by an increase in the ThT signal. After 75 h of incubation, brightfield and fluorescence microscopy reveal the formation of starburst-shaped aggregates (denoted by arrows) and condensates with ThT-positive condensate interfaces. 0.5 µM Kapβ2 inhibits hnRNPA2 amyloid formation while stabilizing the condensed state. Condensates in the presence of Kapβ2 do not exhibit increased ThT fluorescence in their interior or at the interface.

To determine whether Kapβ2 influences hnRNPA2 self-assembly, we analyzed samples containing 4 µM hnRNPA2 (above *c*_sat_) with increasing Kapβ2 concentrations. Sub-stoichiometric Kapβ2 effectively suppressed condensate formation, as visualized by brightfield microscopy (**Fig. 6c**). MRPS analysis confirmed a shift toward smaller assemblies and reduced counts, comparable to controls lacking hnRNPA2 condensates (**Fig. 6d; Suppl. Fig. 11**). Under these conditions, size distributions lost their heavy-tailed character and instead resembled those observed at subsaturated hnRNPA2 concentrations (**Fig. 6b, d**). Parallel experiments with hnRNPA1 yielded similar results, with Kapβ2 strongly inhibiting self-assembly (**Extended Data Fig. 4; Suppl. Figs. 12, 13**). Collectively, these findings extend the mechanism established for FUS and demonstrate that Kapβ2 modulates RBP self-assembly by regulating the pool of free, assembly-competent RBP in the dilute phase, a principle that operates across distinct RBPs with PrLDs. Our findings for hnRNPA2 further support the linked equilibrium model we developed for FUS. Even though hnRNPA2 does not form the types of heavy-tailed distributions we observed for FUS, the effect of Kapβ2 is qualitatively similar. Therefore, the effect of this ATP-independent chaperone is realized primarily via a concentration renormalization effect, whereby the effective concentration of free RBPs is lowered, which affects the overall phase behavior irrespective of the details of the phase behavior.

Next, we examined whether Kapβ2 regulates amyloid formation of hnRNPA2. Previous studies on hnRNPA1 have shown that amyloid formation is initiated within the dilute phase, where primary nucleation and elongation occur, and is further promoted by the condensate interface^46, 49,^_98_. Although interfaces can catalyze fibril formation, likely through secondary nucleation, condensate interiors appear to act as sinks that stabilize against amyloid conversion^49^. Hence, we hypothesized that preferential binding of Kapβ2 to cargo in the dilute phase suppresses amyloid formation by depleting the pool of aggregation-competent monomers.

To test this hypothesis, we examined samples containing 4 µM hnRNPA2 (above the *c*_sat_ for condensate formation) with or without 0.5 µM Kapβ2 (**Fig. 6e**). Thioflavin T (ThT), a fluorescent dye that reports on β-sheet–rich amyloids, was added to monitor fibril formation over time. After 2 hours, brightfield and fluorescence microscopy revealed condensates under both conditions with minimal bulk ThT signal. Over 75 hours, samples lacking Kapβ2 showed a marked increase in ThT fluorescence, consistent with amyloid formation (**Fig. 6e**). Microscopy revealed ThT-positive starburst-like fibrillar structures and condensates with ThT-positive rims, indicating nucleation at condensate interfaces (**Fig. 6e**). By contrast, even after prolonged incubation, samples containing 0.5 µM Kapβ2 maintained consistently low ThT fluorescence and intact condensate morphology without ThT-positive rims, (**Fig. 6e**). These findings demonstrate that low concentrations of Kapβ2 selectively block amyloid formation while preserving condensate integrity, enhancing the metastability of hnRNPA2 condensates. This protection likely arises from preferential binding of

Kapβ2 to hnRNPA2 monomers in the dilute phase, which limits monomer recruitment to condensate interfaces and suppresses fibril nucleation and elongation. Together, our results reveal that Kapβ2 safeguards RBPs from pathological amyloidogenesis by modulating the phase equilibrium and monomer pool.

## Discussion

In this study, we uncovered the mechanism underlying the chaperone activity of the NIR Kapβ2 by dissecting how this NIR regulates self-assembly of the cargo proteins FUS, hnRNPA1, and hnRNPA2. By combining FCS, MRPS, and microscopy, we captured the full spectrum of assemblies from nanometer-scale clusters below *c*_sat_ to micrometer-scale condensates above *c*_sat_. Together, our analyses define how Kapβ2 remodels the dilute phase to prevent aberrant assembly across this entire landscape.

Although both FUS and hnRNPA2 form higher-order assemblies, their self-assembly pathways differ markedly. FUS strongly self-associates even at low nanomolar concentrations, generating a concentration-dependent continuum of clusters that span tens of nanometers to micrometers. These nano- and mesoscale clusters persist below *c*_sat_ and coexist with condensates once *c*_sat_ is breached. This hierarchical assembly behavior was observed not only at the pure protein level but also in cells, where cytoplasmic clusters and condensates form in proportion to FUS expression levels. hnRNPA1 and hnRNPA2 follow a more classical nucleation-driven phase transition^89^, forming few or no clusters below *c*_sat_ and condensing only after surpassing a defined threshold undergoing macrophase separation. This contrasting behavior of hnRNPA1 and hnRNPA2 was reported in recent measurements within germinal vesicles of *Xenopus* oocytes^99^. Condensates composed of wild-type FUS can remain metastable over extended timescales, with solidification initiated at the interface only after several days to weeks^47, 100^. In contrast, hnRNPA2 condensates rapidly convert to amyloids through interface-promoted pathways, consistent with analogous behavior observed for hnRNPA1^46, 98^.

The distinct self-assembly pathways of FUS and hnRNPA2 likely arise from differences in multivalent interaction networks involving their PrLD and arginine-rich regions, which promote phase separation through residue- and domain-specific interaction modes unique to each protein^32,^_51_. For each RBP, FUS and hnRNPA2, we used atomistic simulations to assess how pairs of domains within the protein interact with one another. We also analyzed the patterns of inter-residue distances that underlie the inter-domain interactions (**Extended Data Fig. 5**). This analysis revealed several key differences between the two proteins. In FUS, the normalized distance maps highlight a preference for self-associations between the PrLDs (**Extended Data Fig. 5a**) and a preference for strong associations between the PrLD and the RRM as well as specific sequence stretches within the PrLD and the C-terminal R-rich region, which is broken up into two distinct IDRs (IDR2 and IDR3). Both IDR2 and IDR3 associate strongly (quantified by the inter-residue interactions) with the RRM and the surface of the RRM also enables strong self-associations. These interaction patterns are consistent with previous studies showing that FUS phase separation depends on hydrogen bonding, hydrophobic contacts, and π/sp² interactions spanning a broad range of residues^101–103^. By contrast, the normalized inter-domain, inter-residue distance maps of hnRNPA2 show weaker patterns of association, with the strongest associations involving the PrLD and RRM2 followed by the PrLD and RRM1 and inter-PrLD interactions (**Extended Data Fig. 5b**). The inter-RRM interactions are repulsive, which is in stark contrast to what we observe for FUS, highlighting the differences in the surface-mediated interactions between the RRMs of FUS versus hnRNPA2. The associations of hnRNPA2 associations rely primarily on the PrLD, where evenly spaced aromatic residues (referred to as stickers) and aggregation-prone segments (referred to as zippers) mediate condensation and fibrillization, respectively^46, 57, 104^. Overall, the diversity of association types and strengths that characterize the inter-domain associations of FUS are absent in hnRNPA2, and these distinct interaction patterns may underlie the divergent assembly behaviors of FUS and hnRNPA2, necessitating adaptable chaperone mechanisms such as those provided by Kapβ2.

Kapβ2 provides a unifying regulatory mechanism for higher-order assembly across distinct RBP cargos. Despite the RBP-specific pathways driving phase separation, Kapβ2 controls self-association by selectively engaging monomeric cargo in the dilute phase, thereby renormalizing the pool of free, assembly-competent FUS and hnRNPA2. This coupling between selective recognition and phase equilibria depends on the PY-NLS and enables Kapβ2 to tune condensation across diverse cargos with distinct structures and self-assembly mechanisms. The Kapβ2-to-cargo ratio dictates the concentration of self-association-competent species and thus the resulting assembly state. For example, equimolar Kapβ2:FUS ratios most effectively suppress mesoscopic clustering. By contrast, sub-stoichiometric Kapβ2 suffices to block hnRNPA2 amyloid formation while maintaining condensate integrity.

Regulation is driven by high-affinity, single-site interactions between Kapβ2 and the PY-NLS motifs of cognate cargos. In hnRNPA1 and hnRNPA2, the PY-NLS is embedded within the PrLD, which drives intermolecular interactions essential for self-assembly^5^, whereas in FUS the PY-NLS is positioned outside the PrLD^5^. In all three RBPs, the PY-NLS likely becomes partially buried within dense condensate networks, reducing accessibility to Kapβ2. By contrast, in the dilute phase these motifs remain exposed and available for high-affinity engagement. Through selective sequestration of assembly-competent monomers, Kapβ2 regulates self-assembly across diverse RBPs, independent of intrinsic phase behavior or requirement for ATP hydrolysis. This mechanism enables Kapβ2 to fine-tune phase equilibria, maintain proteome homeostasis, and prevent transitions to pathological aggregation.

Earlier studies showed that NIRs can co-translationally stabilize nascent peptides and shield nuclear proteins, often those containing exposed positively charged regions, from unintended interactions with negatively charged biomolecules in the cytoplasm^105, 106^. These observations suggest that NIRs promote and preserve native protein states and can act in a preventative capacity. Our findings now provide direct evidence linking the binding equilibrium of the Kapβ2–cargo interaction to regulation of the self-assembly behavior of RBPs with PrLDs.

Dysregulated proteostasis and defects in nucleocytoplasmic transport are hallmarks of many neurodegenerative diseases that are closely associated with toxic protein accumulation^65^. In genetic forms of disease, diverse mutations, including those affecting the PY-NLS, can drive cytoplasmic mislocalization of FUS, hnRNPA1, and hnRNPA2, thereby increasing their ability to form fibrils^34, 37, 38^. A prominent example is the FUS^P525L^ variant, in which substitution of proline 525 with leucine stabilizes pre-percolation clusters at low concentrations by two mechanisms: (1) enhancing the intrinsic tendency to form mesoscopic clusters in the dilute phase and (2) weakening Kapβ2 binding, increasing the *K_d_* from ∼50 nM to ∼200 nM^81^. Despite this reduced affinity, Kapβ2 still raises the *c*_sat_ and inhibits condensate formation. Based on our model, a *K_d_* of ∼200 nM predicts a concentration of free FUS^P525L^ of 1.3 µM in the presence of 2 µM Kapβ2. This concentration is below the *c*_sat_ of FUS^P525L^ and thereby results in prevention of condensation under these conditions. The implication is that Kapβ2 efficiently suppresses formation and stabilization of mesoscopic clusters of wild-type FUS but it is far less effective at preventing formation of these assemblies for FUS^P525L^. We suggest that mesoscale clusters act as seeds for amyloid formation in the dilute phase. In individuals carrying the FUS^P525L^ variant, chronic dysregulation of self-assembly combined with cytoplasmic mislocalization likely drives toxic aggregate accumulation and loss of functional assemblies. Cytoplasmic FUS^P525L^ also disrupts the negative autoregulatory feedback loop controlling FUS expression, leading to elevated FUS levels and further cytoplasmic buildup^107^. Previous work has suggested that FUS-linked proteinopathy drives gain-of-toxic-function mechanisms^94, 108^. Dysregulated and persistent clustering could contribute to these mechanisms emphasizing how subtle perturbations of the phase equilibrium could precipitate severe neurodegenerative outcomes.

Our findings demonstrate that Kapβ2 effectively inhibits the interface-promoted, disease-associated conversion of hnRNPA2 condensates into amyloids. Structural studies of Kapβ2 binding to the hnRNPA1 PY-NLS show that this interaction promotes extended cargo conformations that preclude fibril formation^55, 56, 69^, suggesting a similar mechanism operates for Kapβ2 with hnRNPA2. Mechanistically, this inhibitory effect aligns with our model in which Kapβ2 preferentially binds RBP cargo in the dilute phase, stabilizing monomeric states that are incompetent for aggregation.

To date, no disease-associated variants of Kapβ2 have been identified in ALS or FTD, although mutations in other NIRs have been implicated in myopathies and neurodevelopmental disorders^65,^ _109, 110_. Notably, Kapβ2 co-aggregates with FUS in neuronal cytoplasmic inclusions characteristic of FTD-FUS, but not in ALS-FUS^111^. In some cases, hnRNPA1 is also detected within FTD-FUS inclusions^112^. Sequestration of Kapβ2 into these aggregates would reduce the pool of functional NIR and compromise nuclear import and chaperone activity. The resulting depletion could create a self-reinforcing loop that intensifies cytoplasmic mislocalization and aggregation of FUS and hnRNPs, driving proteostatic stress and disease progression.

Our findings outline principles for therapeutic strategies that counter dysregulated self-assembly in disease. Interventions should avoid generating aberrant intermediates and instead stabilize native conformations while preserving access to functional mesoscopic assemblies. Stabilizing the monomeric form alone is unlikely to suffice, and maintaining physiological assembly pathways is essential. The way Kapβ2 binds and regulates monomeric cargo offers a model for therapeutics that preserve native states while supporting proper phase behavior^65^. Reinforcing physiological assembly routes could boost proteostasis capacity to clear toxic aggregates, reduce proteotoxic stress, and slow neurodegenerative progression. This framework provides a foundation for developing strategies that counter protein aggregation diseases.

The mechanism we have uncovered here likely represents a general principle by which ATP-independent chaperones regulate phase behavior to preserve proteome stability. This strategy is likely to extend well beyond NIRs. For example, under physiological conditions, specific short RNA chaperones (25–34 nucleotides) can solubilize FUS and another prion-like RBP, TDP-43^17^, by engaging RNA-binding domains in the dilute phase to allosterically limit PrLD multivalency and restrain aberrant self-association^113–116^. Likewise, TRIM family proteins and polyD/E-rich proteins such as DAXX, which act as potent ATP-independent chaperones against aggregation-prone substrates, may harness related principles to suppress pathological phase transitions and maintain proteome organization^117–120^. While control of self-assembly by remodeling the dilute phase is likely to be a major regulatory mode, these ATP-independent chaperones may also directly remodel preassembled states by harnessing binding energy to disrupt intermolecular contacts via mechanisms which remain to be fully elucidated^60, 118–124^. Together, these findings define a unifying paradigm in which ATP-independent chaperones sustain proteome homeostasis by controlling the dynamic balance between soluble proteins, reversible assemblies, and pathological aggregates.

## Methods

### Cloning

All constructs for recombinant protein expression and purification were designed as MBP fusion constructs, with an interspersed TEV protease cleavage site. hnRNPA1 and hnRNPA2 also contain a C-terminal 6x His tags. MBP-hnRNPA1-6xHis and MBP-hnRNPA2-6xHis and MBP-FUS constructs were cloned into the pMAL-C2T plasmid background. Similarly, Kapβ2 was cloned into the pE-His-SUMOpro vector. For experiments in HEK293T cells, GFP-FUS-NES, mCherry-Kapβ2, and the CluMPS reporter sequences were cloned into the pCMV plasmid background. All sequences were verified via full plasmid sequencing (Plasmidsaurus).

### Protein purification

MBP-hnRNPA1-6xHis and MBP-hnRNPA2-6xHis were transformed into BL21 DE3 RIL Escherichia coli (E. coli) cells via heat shock (35 s at 42 °C) and plated on LB agar plates supplied with 100 µg/ml ampicillin. After incubation overnight at 37 °C, 5 ml LB pre-cultures were inoculated and grown overnight at 37 °C. The next day, cultures were supplied with 50 % glycerol, flash-frozen in liquid nitrogen and stored at -80 °C for further use. For scale-up, glycerol stocks were used to inoculate 5 ml pre-cultures in LB media containing 100 µg/ml ampicillin. The MBP-FUS construct was transformed into One Shot BL21 Star (DE3) chemically competent Escherichia coli cells (Thermo Fisher Scientific) via heat shock and plated onto LB amp agar plates containing 100 µg/ml ampicillin. pE-SUMO-pro plasmids containing the His-SUMO-Kapβ2 construct^73^ were transformed into BL21 DE3 RIL cells as described above.

Cultures were scaled up by using pre-cultures (hnRNPA1 and hnRNPA2) or by scraping off colonies from agar plates to inoculate 1L LB liquid media containing 100 µg/ml ampicillin and 0.02 % glucose. To induce protein expression, 1 mM IPTG (FUS, Kapβ2) or 0.5 mM IPTG (hnRNPA1, hnRNPA2) were added to the cultures when reaching an optical density (OD) of ∼ 0.5. Protein expression was carried out over night for 16-18 h at 16 °C. Cells were harvested via centrifugation (4000 RPM, 20 min, 4 °C). Cell pellets were flash-frozen in liquid nitrogen and stored at -80 °C until further use.

hnRNPA1 and hnRNPA2 were purified by resuspending cell pellets in resuspension buffer (20 mM HEPES, pH 7.5, 50 mM NaCl, 2 mM EDTA, 10 % glycerol, 2 mM DTT, 10 µg/ml DNase, 10 µg/ml RNase, cOmplete EDTA-free protease inhibitor tablets (Roche)). After resuspension, 50 µg/ml lysozyme were added, followed by incubation for 30 min on ice. After lysis via sonication, lysates were centrifuged (20 000 RPM, 45 min, 4 °C) to separate soluble from insoluble pellet fraction. The soluble supernatant was combined with 5 ml amylose resin (New England Biolabs) equilibrated in resuspension buffer and incubated for 2h at 4 °C. Afterwards, beads were washed with resuspension buffer and bound proteins were eluted with Amylose Elution Buffer (20 mM HEPES, pH 7.5, 50 mM NaCl, 2 mM EDTA, 10 % glycerol, 10 mM maltose). To remove RNA, the eluted sample was loaded onto a Heparin column (HiTrap^TM^, Heparin HP, Cytiva) equilibrated in Buffer A (same composition as Amylose Elution Buffer) using an FPLC system (ÄktaPurifier, GE Healthcare). Bound proteins were eluted via a salt gradient using high-salt Buffer B (20 mM HEPES, pH 7.5, 1 M NaCl, 2 mM EDTA, 10 % glycerol, 2 mM DTT). hnRNPA1 fractions of interest elute at ∼ 40 % Buffer B, hnRNPA2 fractions elute at ∼ 30 % Buffer B. Fractions containing the protein of interest were collected and concentrated for further polishing via Size Exclusion Chromatography (SEC) on a 16/600 Superdex 200 column (Cytiva) using SEC buffer (20 mM HEPES, pH 7.5, 200 mM NaCl, 2 mM EDTA, 10 % glycerol, 2 mM DTT).

MBP-FUS purification was performed as previously described^73^. In brief, pellets were thawed on ice and resuspended in lysis buffer (20 mM HEPES, pH 7.4, 50 mM NaCl, 2 mM EDTA, 10 % glycerol, 2 mM DTT, cOmplete EDTA-free protease inhibitor tablets (Roche)). Resuspended pellets were supplied with 50 µg/ml lysozyme and incubated on ice for 30 min and lysed via sonication. After centrifugation at 16 000 RPM for 20 min at 4 °C (Avanti J-E centrifuge, Beckman Coulter) using a JA-20 rotor (Beckman Coulter), the soluble supernatant was combined with 5 ml amylose resin (New England Biolabs) and incubated for 2h at 4 °C. Amylose beads were washed with lysis buffer and bound proteins were eluted using elution buffer (20 mM HEPES, pH 7.4, 50 mM NaCl, 2 mM EDTA, 10 % glycerol, 2 mM DTT, 10 mM maltose). To remove bound RNA, the eluted sample was loaded on a 5 ml Heparin column (HiTrap^TM^, Heparin HP, Cytiva) equilibrated in Buffer A (20 mM HEPES, pH 7.4, 50 mM NaCl, 2 mM EDTA, 10 % glycerol, 2 mM DTT) using an FPLC system (ÄktaPurifier, GE Healthcare), washed with Buffer A and eluted via a gradient using Buffer B (20 mM HEPES, pH 7.4, 1 M NaCl, 2 mM EDTA, 10 % glycerol, 2 mM DTT). Protein-containing fractions were analyzed via SDS-PAGE followed by Coomassie staining. Pure fractions elute at ∼ 60 % Buffer B (20 mM HEPES, pH 7.4, 620 mM NaCl, 2 mM EDTA, 10 % glycerol, 2 mM DTT) and were pooled, concentrated and flash-frozen in liquid nitrogen.

For His-SUMO-Kapβ2 purification, pellets were resuspended in lysis buffer (50 mM Tris, pH 7.4, 100 mM NaCl, 20% glycerol, 10 mM imidazole, pH 8, 2.5 mM β-mercaptoethanol, cOmplete EDTA-free protease inhibitor tablets (Roche)) and lysed via sonication. For batch Ni-NTA affinity chromatography, lysed cells were centrifuged (20 min, 16 000 RPM, 4 °C) and the supernatant was incubated with 5 ml Nickel NTA agarose beads (Qiagen) for 75 min at 4 °C. Beads were washed with lysis buffer and proteins were eluted using elution buffer (50 mM Tris, pH 7.4, 100 mM NaCl, 20% glycerol, 200 mM imidazole, pH 8, 2.5 mM β-mercaptoethanol). Eluted samples were exchanged into Buffer A (20 mM imidazole, pH 6.5, 75 mM NaCl, 20 % glycerol, 2 mM DTT) using Amicon® Ultra-15 Centrifugal Filters (MWCO 100,000 Da, Merck Millipore). Ulp1 was added in a 1:100 molar ratio and cleavage of the His-SUMO tag was carried out overnight at 30 °C. The next day, cleaved samples were further purified using anion exchange chromatography (HiTrap^TM^ Q HP, Cytiva). Samples were eluted via gradient elution using Buffer B (20 mM imidazole, pH 6.5, 1 M NaCl, 1 mM EDTA, 20 % glycerol) using an FPLC system. All collected fractions were analyzed via SDS-PAGE and Coomassie staining. Pure fractions eluted at 20 % Buffer B.

To purify TEV protease, frozen pellets were thawed and resuspended in lysis buffer (25 mM Tris, pH 8.0, 500 mM NaCl, 1 mM DTT, cOmplete EDTA-free protease inhibitor tablets (Roche)). Cell lysis was conducted via sonication, followed by centrifugation (20 000 RPM, 4 °C, 30 min) to separate soluble from insoluble pellet fraction. The supernatant was mixed with 5 ml Ni-NTA resin, equilibrated with lysis buffer. After incubation for 1h at 4 °C, the beads were washed with Wash Buffer (25 mM Tris, pH 8.0, 500 mM NaCl, 1 mM DTT, 25 mM imidazole, pH 8) and eluted using Elution Buffer supplied with high concentrations of imidazole (25 mM Tris, pH 8, 500 mM NaCl, 1 mM DTT, 300 mM imidazole, pH 8.0). The eluted sample was further purified via SEC on a 16/600 Superdex 200 column (Cytiva) using SEC buffer (25 mM Tris, pH 7.0, 300 mM NaCl, 10 % glycerol, 1 mM DTT).

For all purifications, all collected fractions were analyzed via SDS-PAGE and Coomassie staining after each purification step. The final, pure fractions were pooled, concentrated, flash-frozen in liquid nitrogen, and stored at -80 °C. Representative chromatograms and SDS-PAGE analysis are shown in **Suppl. Fig. 14**.

### Protein self-assembly assays

Phase separation of all constructs was induced by in-situ cleavage of the MBP solubility tag by TEV protease. MBP-FUS, MBP-hnRNPA1 and MBP-hnRNPA2 were diluted to 3x the final assay concentration in their respective elution buffers. Similarly, TEV protease and Kapβ2 were diluted to 3x the final assay concentration in phase separation buffer (20 mM HEPES, pH 7.4, 1 mM DTT). For Thioflavin T (ThT) kinetics, the phase separation buffer was supplied with 20 µM ThT. To obtain the desired conditions, 10 µl of each component were mixed in 384-well plates (Matriplate, Azenta). The final assay buffer conditions for FUS are 20 mM HEPES, pH 7.4, 207 mM NaCl, 1.3 mM DTT, 0.7 mM EDTA, 3.3 % glycerol. For hnRNPA1 and hnRNPA2, the final assay buffer conditions are 20 mM HEPES, pH 7.4, 66.7 mM NaCl, 1.3 mM DTT, 0.7 mM EDTA, 3.3 % glycerol. FUS, hnRNPA1 and hnRNPA2 protein concentrations are indicated in the figures, the concentration of TEV protease is 0.04 mg/ml in all assays. Samples were incubated at room temperature for 1.5-2 h, to ensure full cleavage of the MBP tag by TEV protease inducing protein self-assembly. Widefield and fluorescence microscopy were performed on an EVOS M-5000 microscope (Thermofisher Scientific). Thioflavin T kinetic assays were performed on a ClarioStar plate reader (BMG Labtech) (excitation wavelength 450 nm / emission wavelength 490 nm).

### Microfluidic Resistive Pulse Sensing (MRPS)

Microfluidic Resistive Pulse Sensing (MRPS) was performed on an nCS1 instrument (Spectradyne). Particle sizes are measured by injecting 5 µl of a sample of interest into a microfluidic cartridge, provided by the manufacturer. Samples were prepared as described above, in Lo-bind Eppendorf tubes to minimize sticking to test tube walls. To measure size distributions covering higher-order structures across several orders of magnitude, two different cartridges with different were used. C-400 cartridges capture assemblies from 70-400 nm, C-2000 cartridges measure assembly sizes between 250-2000 nm.

Each condition was measured in triplicates with each cartridge. To accurately combine the size distributions from both cartridges to full-range distributions (70-2000 nm), we generated synthetic datasets by randomly selecting subsets of values measured from the replicates. These data sets were accumulated, obtaining means and spread of the combined distributions.

## Fluorescence Correlation Spectroscopy (FCS)

For Fluorescence Correlation Spectroscopy (FCS), samples were prepared as described above. To track fluorescence intensity over time, 16 – 100 % of MBP-FUS were replaced with fluorescein-labeled MBP-FUS (MBP-FAM-FUS). FCS measurements were performed on a confocal microscope (Confocor II LSM, Zeiss) with a 40x water immersion objective. For each sample, 50 x 10s fluorescence intensity traces were collected and analyzed by using the Confocor II FCS software (Zeiss), correlating the fluorescence intensity at time point t to the intensity at time point t+*τ*. To extract the characteristic diffusion time *τ_D_* of each trace, equation 1 was fit to the extracted autocorrelation function G(t), assuming a single-component system.

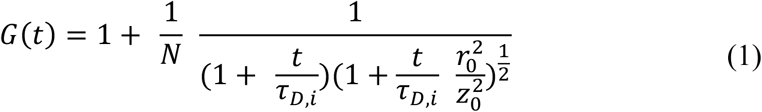

Here,*τ_D_* is the diffusion time, *N* is the number of fluorophores in the confocal detection volume, t is the correlation time, r*_0_* is the resolution in the x-y plane, and *z_0_* is the resolution in the z plane.

To extract hydrodynamic radii *R_h_*, the Stokes-Einstein equation was applied (eq. 2 and 3)

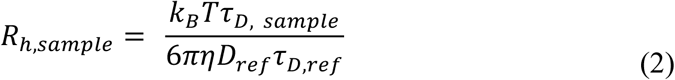

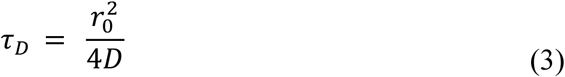

where *k_B_* is Boltzmann’s constant, *T* is the temperature in Kelvin, *η* is the viscosity of the buffer, and *D* is the diffusion coefficient.

### Comparison of size distributions via Kullback-Leibler (KL) divergence

We calculated KL divergences to interrogate how two distributions are different from each other. To avoid computational errors, we set all zeros arbitrarily to 1*10^-5^. We then converted all size distributions into probability density functions. Then we computed KL divergence values (*KLD*) using the following equation (eq. 4):

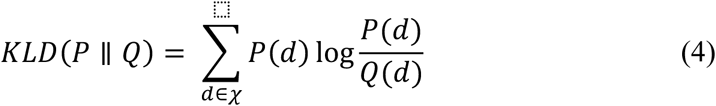

where *d* is the particle diameter, *P(d)* is the probability density function of a sample of 2 µM FUS at increasing concentrations of Kapý2, and *Q(d)* is the probability density function of a sample containing only FUS.

### Model

Previous extensive analysis of the Kapβ2-FUS interaction has shown that the binding of the two proteins is mediated by a high-affinity interaction via the C-terminal PY-NLS. K_D_s have been measured with Isothermal Titration Calorimetry (ITC) and fluorescence anisotropy, and reported to lie in the range of 10 – 100 nM^61, 70, 80, 81^.

We assume a simple binding equilibrium, based on a single interaction site between FUS and Kapβ2 (eq. 5).

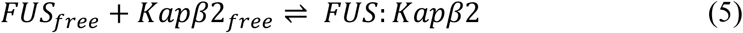

This leads to the following expression of the dissociation constant K_D_ (eq. 6):

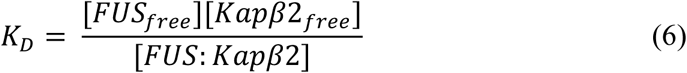

To extract the concentration of free FUS [*FUS_free_*] depending on the total concentration of Kapβ2 [*Kapβ2_tot_*], we assume (eqs. 7 and 8):

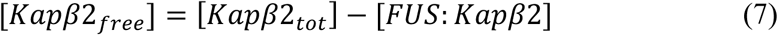

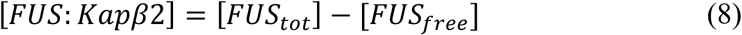

At a given total concentration of FUS [*FUS_tot_*] and dissociation constant K_D_ for the FUS: Kapβ2 complex, this allows us to calculate the concentration of [*FUS_free_*] as a function of the total Kapβ2 concentration [*Kapβ2_tot_*] (**Fig. 4**).

### Cell culture and plasmid transfection

Lenti X HEK 293T (Takara Bio) were maintained in standard cell culture incubators at 37°C and 5% CO_2_. Cells were cultured in DMEM containing 10% fetal bovine serum (FBS) and 1% penicillin/streptomycin (P/S). For live cell imaging experiments, cells were seeded in 384 well plates (Cellvis) with 4000 cells and 50μL media per well. 384 well plates were coated with 10 μg/mL Human Plasma Fibronectin Purified Protein (Sigma-Aldrich) diluted in PBS 1 h before seeding. Plates were centrifuged at 20 x g for 1 min to settle cells immediately after addition of cells. Cells were transfected 24 h after seeding using Lipofectamine™ 3000 Transfection Reagent (ThermoFisher Scientific) with each well receiving a final transfection mix of 2.5-50ng plasmid DNA, 0.05 μl Lipofectamine™ reagent, 0.05 μL P3000 reagent, and brought to total volume of 2.5 μl with Opti-MEM medium (ThermoFisher Scientific). Wells were ‘poly-transfected’ with each plasmid to achieve largely de-correlated expression patterns^125^. Cell media was changed 24 h after transfection.

### Live-cell imaging

Cells were imaged 48 hours after transfection, and 5 μg/ml Hoechst 33342 (Sigma-Aldrich) was added 20 min prior to imaging. Live-cell imaging was performed using a Nikon Ti2-E microscope equipped with a Yokagawa CSU-W1 spinning disk, 405/488/561/640 nm laser lines, an sCMOS camera (Photometrics), 40X air objective, and a motorized stage equipped with an environmental chamber that maintained temperature at 37°C and 5% CO_2_.

### Cell segmentation and condensate quantification

Cell body segmentation and nuclear segmentation was performed using Cellpose^126^. FUS condensates were segmented using a previously developed MATLAB algorithm for identifying condensate pixels from images with cytoplasmic segmentations^92^. Condensate data visualization was generated in RStudio with the tidyR packages^127^.

### Data analysis

Condensate areas were extracted using in-house written code in MATLAB (version R2020a). Micrographs are representative images which were adjusted in Fiji (ImageJ2, version 2.3.0/1.53q) to visualize the condensates. MRPS size distributions were extracted in the Spectradyne Analysis Software (Spectradyne Tools, version 3.6.0.9) and plotted using MATLAB. FCS autocorrelation functions and diffusion times *τ_D_* were extracted using the Zeiss Confocor II FCS software, hydrodynamic radii were calculated and plotted in MATLAB.

### Atomistic simulations

All-atom Metropolis Monte Carlo (MC) simulations were performed using the ABSINTH implicit solvent model and forcefield paradigm as made available in the CAMPARI simulation package (http://campari.sourceforge.net)^128^. The simulations utilized the abs_3.5_opls.prm parameter set in conjunction with optimized parameters for neutralizing and excess Na+ and Cl-ions^129^. In each simulation, we used spherical droplets with a radius of 200 Å to accommodate pairs of interacting domains. The salt concentration was 1.7 mM NaCl. For each protein, we excised pairs of domains X and Y and performed simulations of freely diffusing and associating domains. In detail, each simulation comprised two domains X and Y that were either a pair of folded domains (typically RRMs), a pair of IDRs (PrLDs or R-rich IDRs), or an RRM and an IDR. The approach we followed was that of Shinn et al.^99^. From the simulations, we extracted statistics for inter-residue distances using SOURSOP^130^. These statistics were used to compute the ensemble-averaged distances 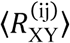 between pairs of residues *i* and *j*, where the former is located on domain X and the latter on domain Y. The observed pattern of inter-domain, inter-residue distances were normalized using priors obtained for non-interacting pairs of the domains as described by the Shinn et al.^99^. The data are shown as normalized and properly referenced inter-residue, inter-domain interaction coefficients 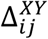. These coefficients quantify the effective attractions or repulsions between pairs of residues, *i* and *j* across a pair of domains X and Y. Note that 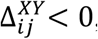, implies inter-residue attractions, 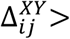 0 , implies inter-residue repulsions, and 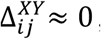, implies that the pair of residues are non-interacting or ideal. Also, the values of the interaction coefficients are bounded such that 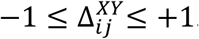

## Acknowledgements

We are grateful for the support of Dr. Luca Musante (Extracellular Vesicle Core, UPenn) for assistance with MRPS measurements. The original measurements were performed using the MRPS nCS1 instrument in the Center for Biomolecular Condensates at Washington University in St. Louis. We also thank Johanna Bergstrom for pilot experiments, Dr. Lijun Zhou and Lisa Nakayama for providing FAM-labeled FUS protein, and Dr. Bede Portz for providing glycerol stocks to express hnRNPA1 and hnRNPA2 constructs. We thank all members of the Shorter Lab for helpful discussions and feedback on the project. We thank Dr. Edward Barbieri, Dr. Charlotte Fare, Dr. Raju Roy, Dr. Hana Odeh, and Zarin Tabassum for helpful comments on the manuscript. This work was supported by a Mildred Cohn Distinguished Postdoctoral Award (M.L.), an ALS Association Milton Safenowitz Postdoctoral Fellowship (M.L.), an Alzheimer’s Association Research Fellowship (M.L.), an ALS Scholars in Therapeutics Award (Sean M. Healey & AMG Center for ALS, Find MND, ALS Finding a Cure, M.L.), Target ALS (J.S.), the ALS Association (J.S.), the Office of the Assistant Secretary of Defense for Health Affairs through the Amyotrophic Lateral Sclerosis Research Program W81XWH-20-1-0242 (J.S.), the G. Harold & Leila Y. Mathers Foundation (J.S.), and NIH grants R01AG077771 (J.S.), R01NS121114 (R.V.P.), K99GM152778 (M-K.S.), and R35GM138211 (L.J.B.).

## Contributions

Conceptualization, M.L., R.V.P., & J.S.; methodology, M.L., M.K.S., R.V.P., & J.S.; validation, M.L., M.K.S., & T.R.M..; formal analysis, M.L., M.K.S., T.R.M., V.L., L.J.B. R.V.P., & J.S.; investigation, M.L., M.K.S., & T.R.M.; resources, M.L., M.K.S., R.V.P., & J.S.; data curation, M.L., M.K.S., T.R.M., V.L., R.V.P., & J.S.; writing – original draft, M.L., & J.S.; writing – review & editing, M.L., M.K.S., T.R.M., L.J.B., V.L., R.V.P., & J.S.; visualization, M.L., M.K.S., T.R.M., V.L., L. J.B., R.V.P., & J.S.; supervision, M.L., M.K.S., L.J.B., R.V.P., & J.S.; project administration, M.L., R.V.P., & J.S.; funding acquisition, M.L., M.K.S., L.J.B., R.V.P., & J.S.

## Data and code availability

All relevant data are included in the main, extended and supplementary figures. Source data will be provided with the manuscript. Codes to analyze and plot micrographs, MRPS and FCS data will be uploaded to a publicly accessible server. All plasmids and reagents are available from the corresponding authors.

---

**Supplementary Fig. 1.**
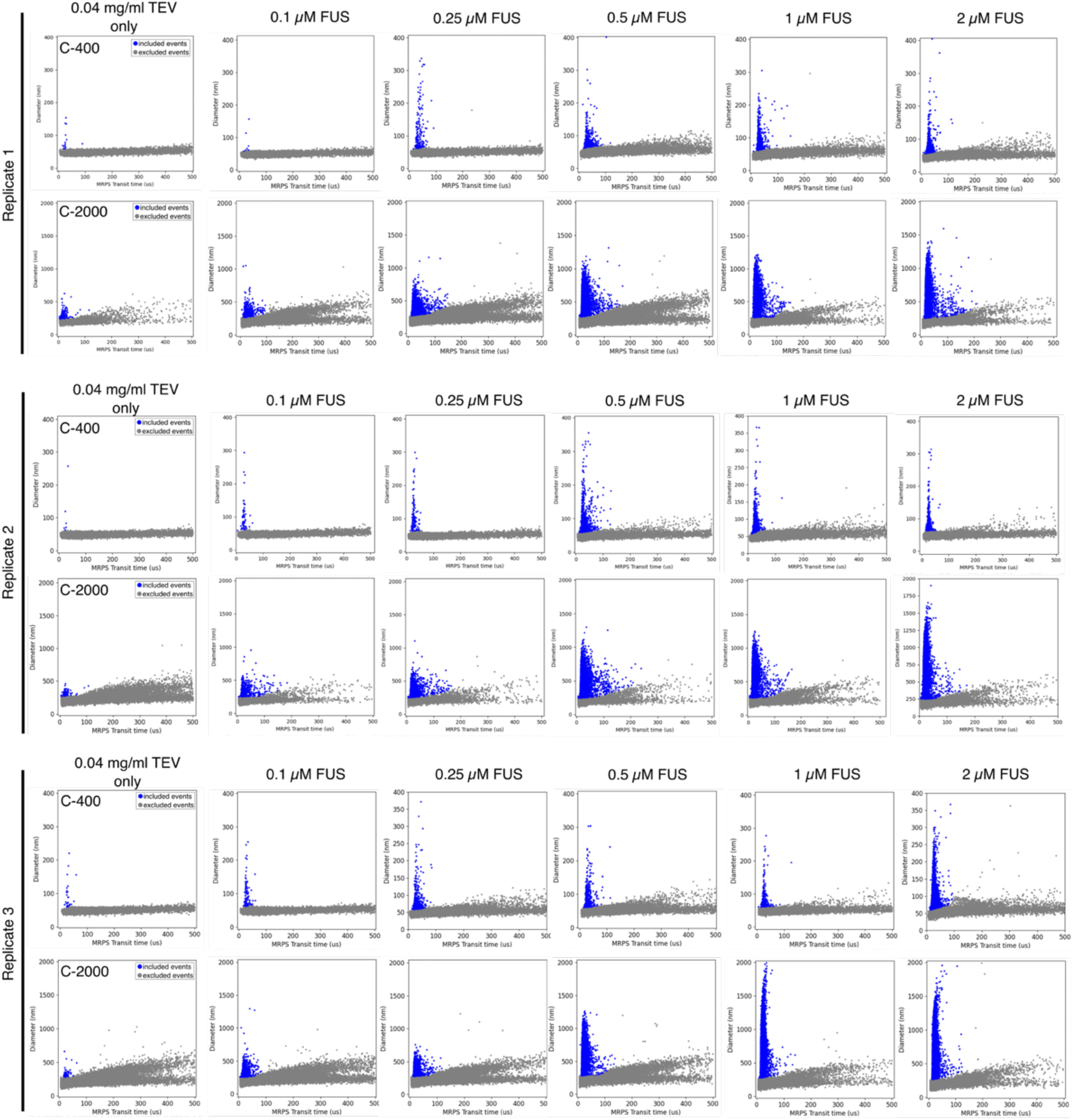
Microfluidic Resistive Pulse Sensing (MRPS) scatter plots of samples at increasing FUS concentrations. Three independent replicates of each condition were measured using two types of cartridges with different nano constriction widths (C-400 and C-2000). 0.04 mg/ml TEV protease was measured as control. Scatter plots show the measured particle diameter as a function of the MRPS transit time. Each blue dot represents a single event, corresponding to a single particle crossing the nano constriction. These scatter plots were used to extract size distributions shown in Fig. 2b. Gray dots represent the noise floor of the instrument and were excluded from the analysis.

**Supplementary Fig. 2:**
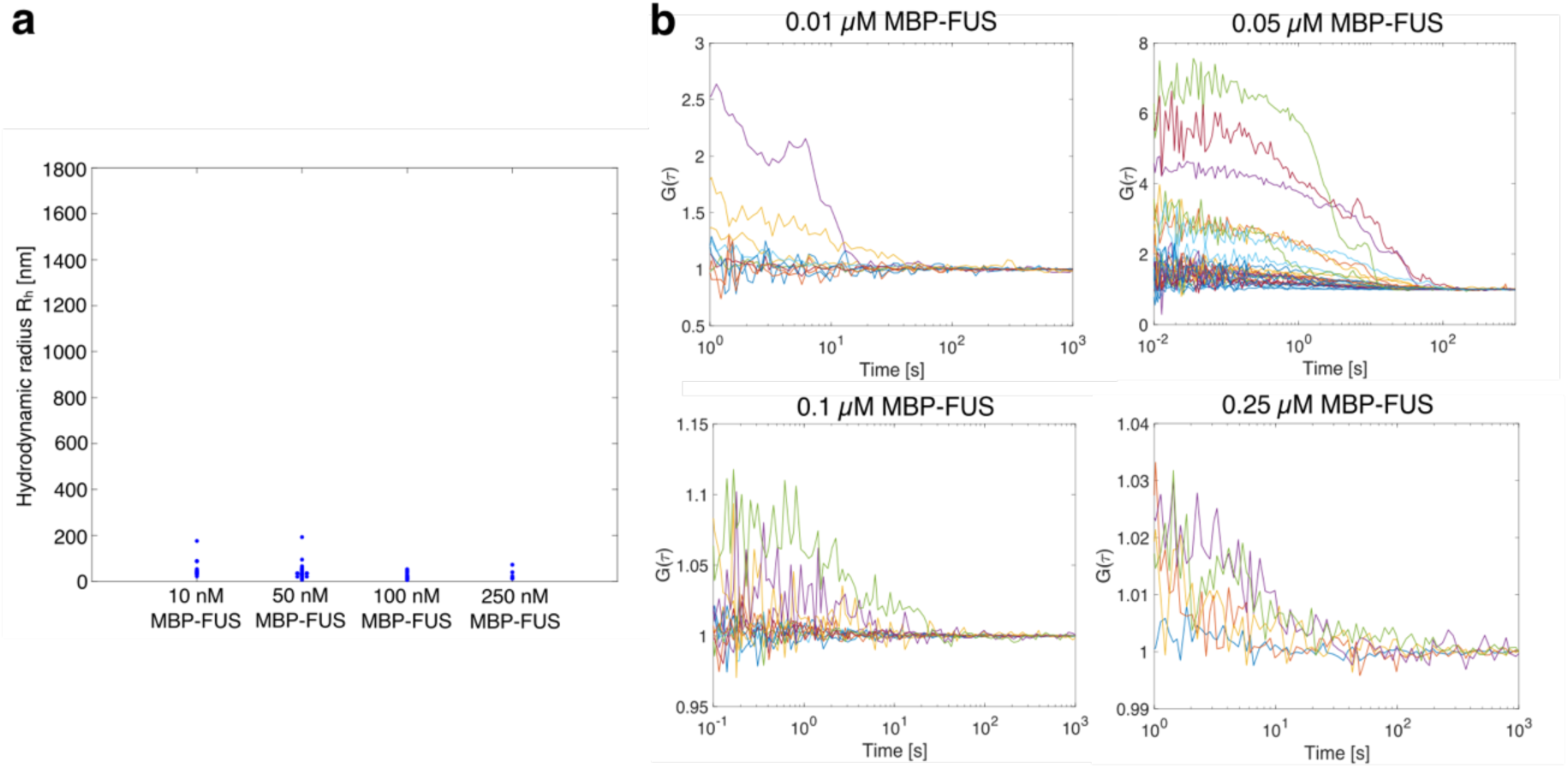
Fluorescence Correlation Spectroscopy (FCS) of MBP-FUS in absence of TEV protease. **a**, Without TEV protease, MBP-FUS particle sizes remain largely below 100 nm. **b,** FCS autocorrelation functions of samples in a.

**Supplementary Fig. 3:**
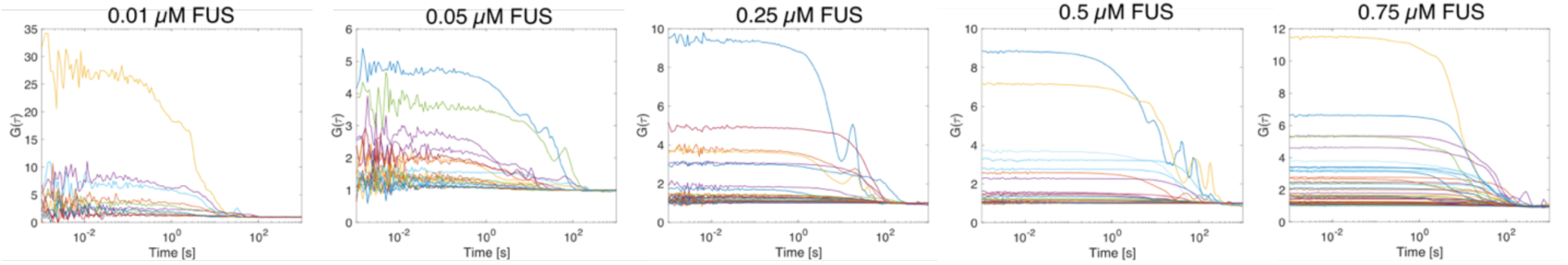
FCS autocorrelation functions of samples at increasing FUS concentrations. Autocorrelation functions were extracted from fluorescence intensity fluctuations over 10 s. A single-component model was fitted to determine characteristic diffusion times 1_D_ and particle hydrodynamic radii R_h_. Particle sizes are showed in Fig. 2c.

**Supplementary Fig. 4:**
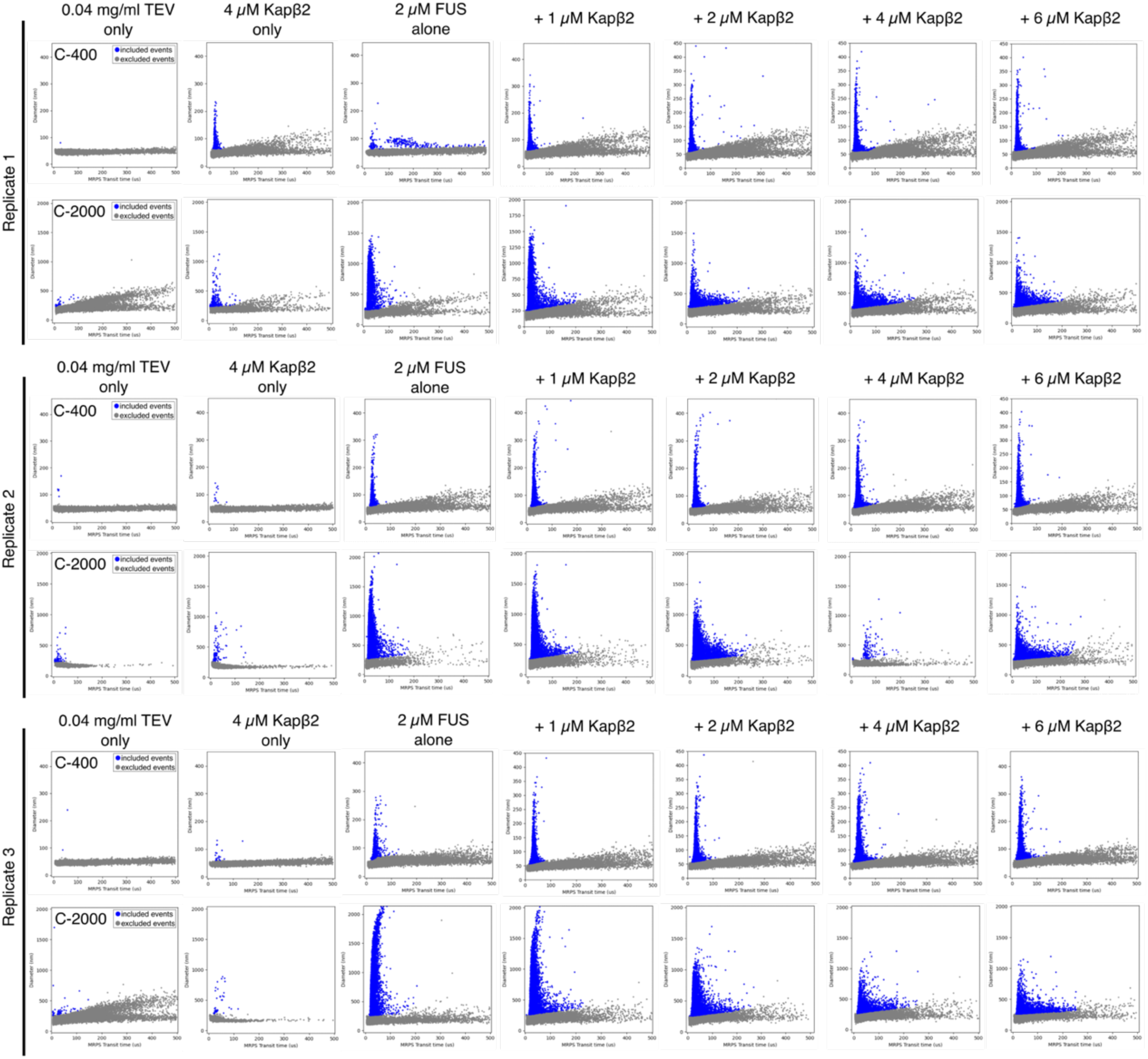
Microfluidic Resistive Pulse Sensing (MRPS) size scatter plots of samples at 2 µM FUS in absence and presence of increasing Kapβ2 concentrations. 0.04 mg/ml TEV protease and 4 µM Kapβ2 alone were analyzed as control samples. Each condition was measured as three independent replicates with two types of cartridges (C-400 and C-2000). Scatter plots show the measured particle diameter as a function of the MRPS transit time. Blue dots represent single assemblies which were used to extract size distributions depicted in Fig. 2d. Gray dots correspond to the noise floor of the instrument and were excluded from the analysis.

**Supplementary Fig. 5:**
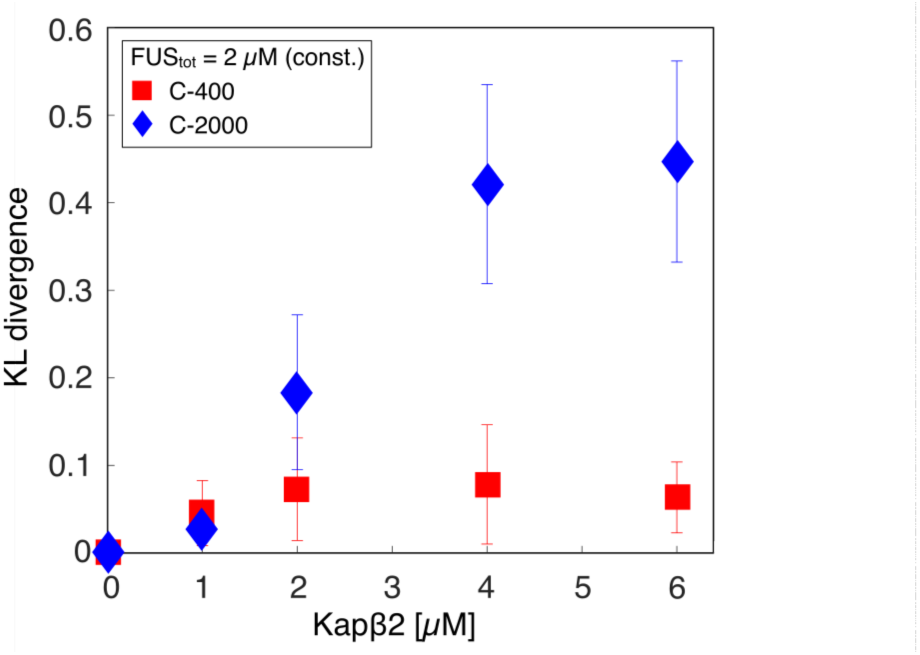
Kullback-Leibler (KL) divergence of size distributions of FUS in presence of Kapβ2 with respect to FUS alone. The KL divergence increases with increasing Kapβ2 concentrations, plateauing at ≥ 4 µM Kapβ2. Mean KL divergence was obtained from size distributions measured as triplicates using C-400 and C-2000 cartridges. Error bars represent the standard deviation.

**Supplementary Fig. 6:**
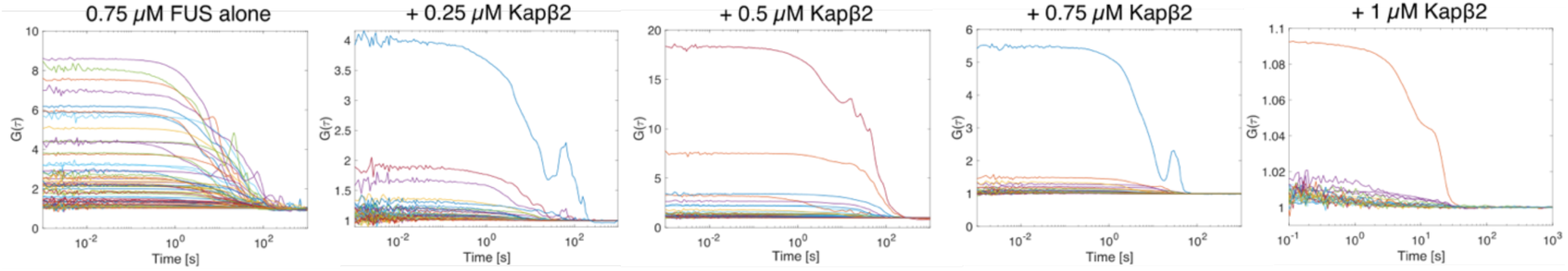
FCS autocorrelation functions of FUS samples in absence and presence of increasing Kapβ2 concentrations. Autocorrelation functions were extracted from fluorescence intensity fluctuations over 10 seconds. A single-component model was used to extract characteristic diffusion times 1_D_ to compute particle hydrodynamic radii R_h_ which are shown in Fig. 2f.

**Supplementary Fig. 7:**
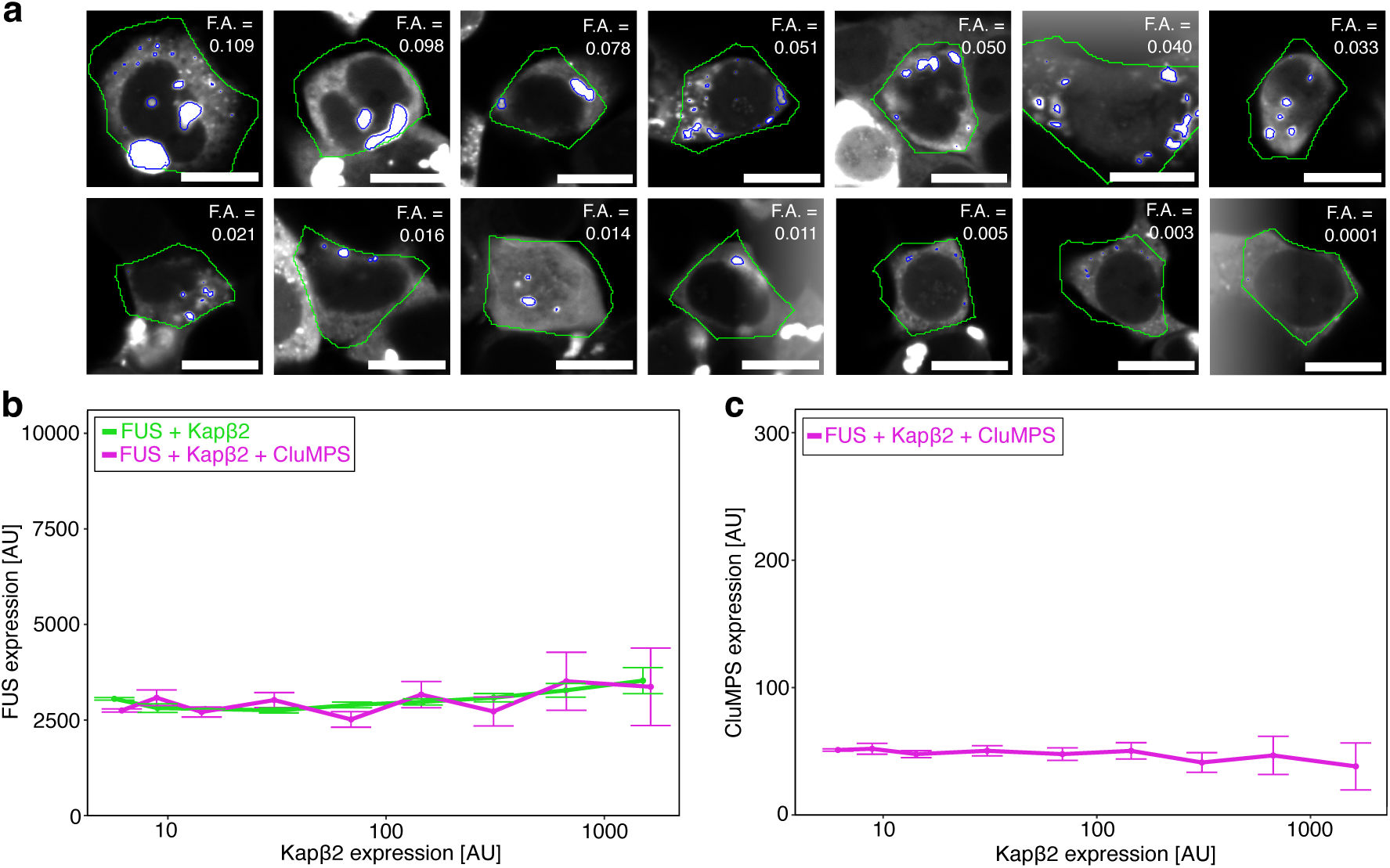
Expression levels of FUS and the CluMPS reporter across increasing Kapβ2 concentrations. **a,** Representative images of cells exhibiting increasing extent of FUS assembly, as quantified by extracting the fraction of condensate area / cell area (F.A.). Scale bars, 20 µm. **b,** Effect of increasing expression levels of Kapβ2 on FUS condensation is not driven by FUS expression levels. **c,** Effect of increasing expression levels of Kapβ2 on FUS condensation is not driven by expression levels of the CluMPS reporter.

**Supplementary Fig. 8:**
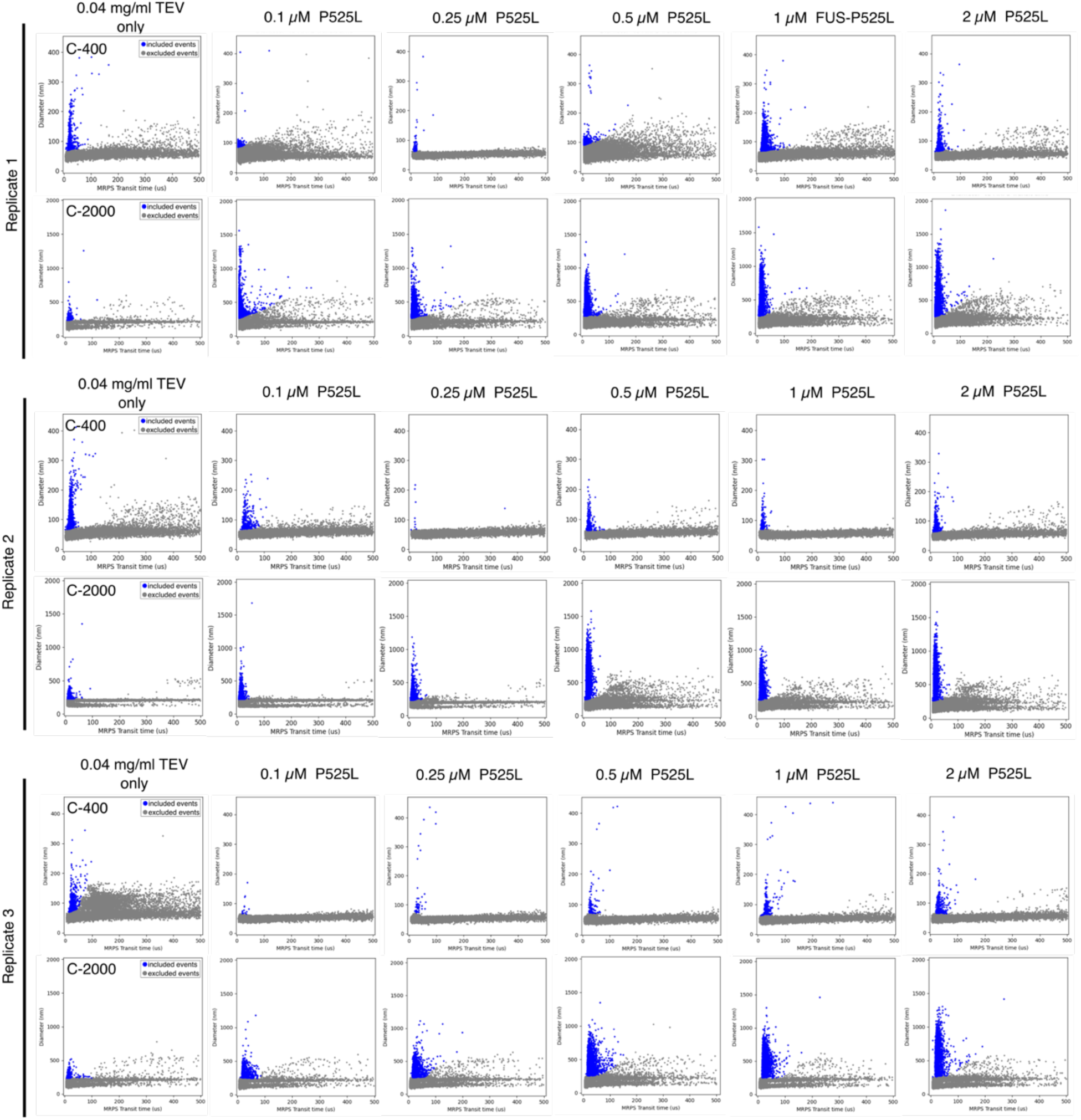
Raw MRPS size scatter plots of samples at increasing FUS^P525L^ concentrations. Scatter plots depict the diameters of single assemblies as a function of the MRPS transit time. Data correspond to size distributions in Fig. 5d. 0.04 mg/ml TEV protease served as control sample. Each blue dot is the diameter of a single particle, gray dots represent the noise floor of the instrument and were excluded. All samples were measured using two types of cartridges with different nano constriction widths (C-400 and C-2000) and replicated three times.

**Supplementary Fig. 9:**
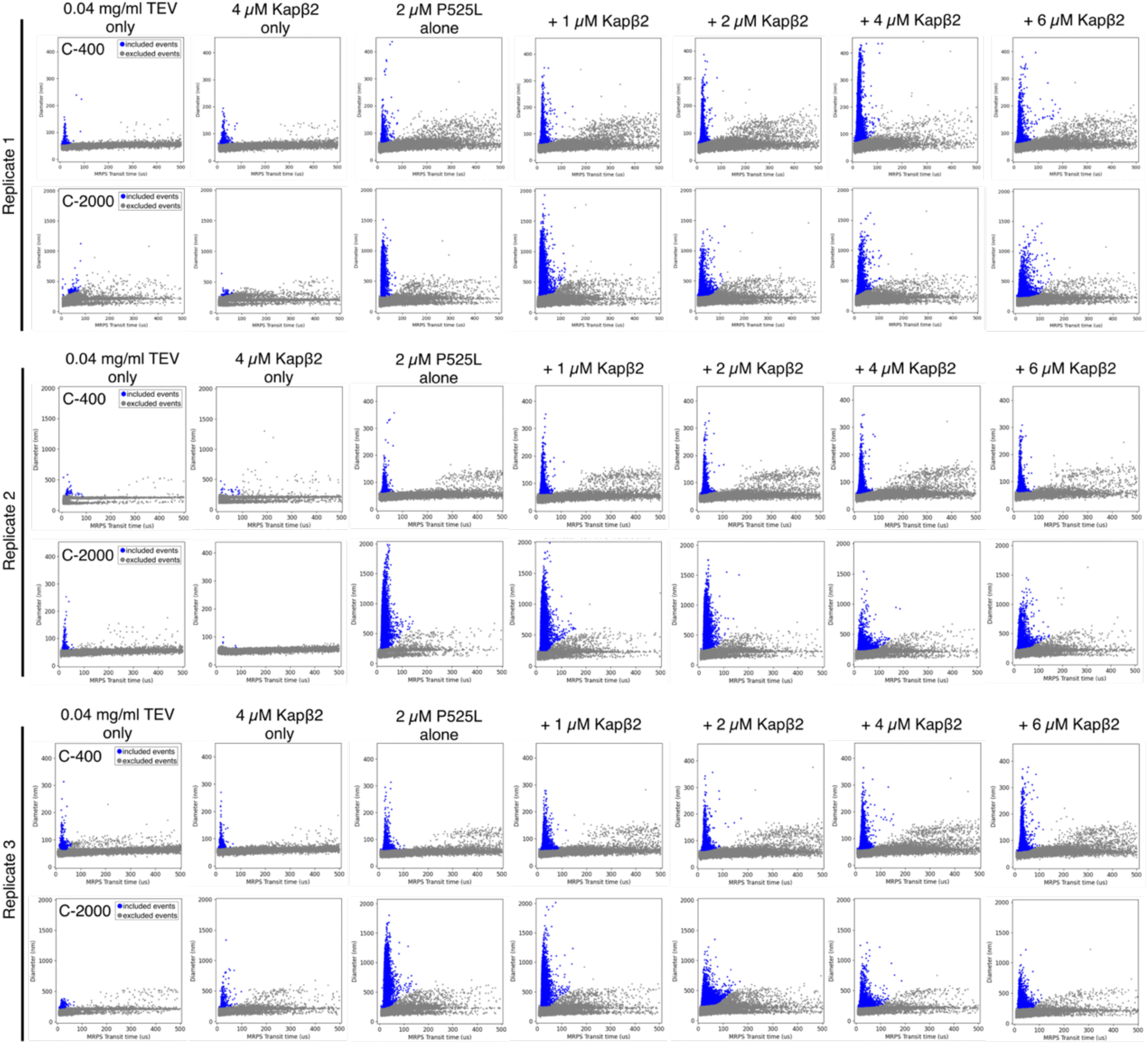
Raw MRPS scatter plots of FUS^P525L^ samples in absence and presence of increasing concentrations of Kapβ2. Plots show three replicates of the particle diameter as a function of the MRPS transit time. Each sample was measured using cartridges with a narrower (C-400) and a wider (C-2000) nano constrictions. 0.04 mg/ml TEV protease and 4 µM Kapβ2 alone were analyzed as control samples. Each blue dot represents a single particle detected. Gray dots correspond to the noise floor of the instrument and were not counted as events. Size distributions extracted from this data are shown in Fig. 5f.

**Supplementary Fig. 10:**
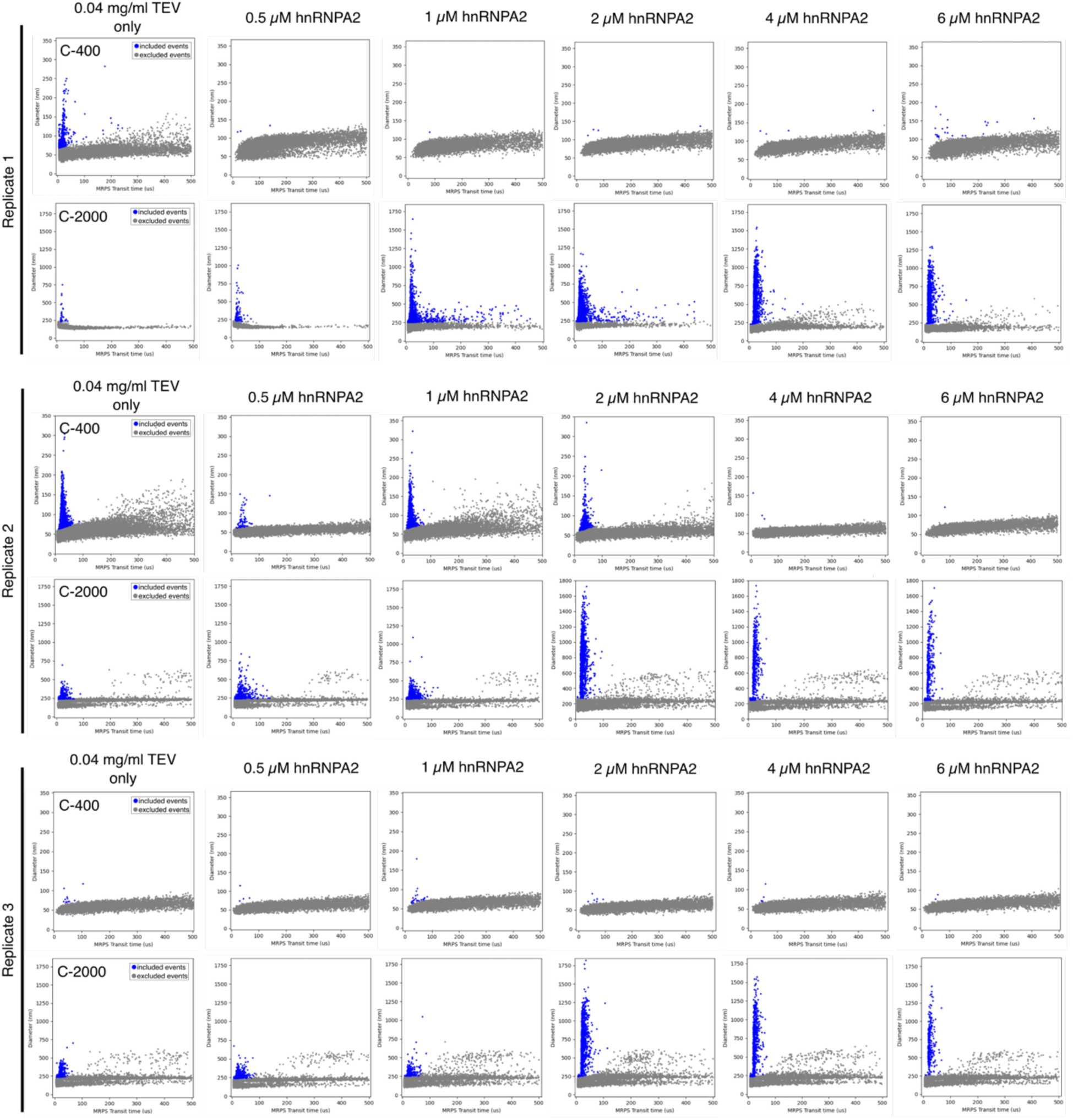
Raw MRPS scatter plots of samples at increasing hnRNPA2 concentrations. Plots show the particle diameter dependent on the MRPS transit time. Each blue dot is a single particle crossing the nano constriction. All samples were measured in triplicates using two different cartridges (C-400 and C-2000) to capture a larger range of particle sizes. 0.04 mg/ml TEV protease was measured as control. Gray dots represent the noise floor. Scatter plots were used to extract particle size distributions depicted in Fig. 6b.

**Supplementary Fig. 11:**
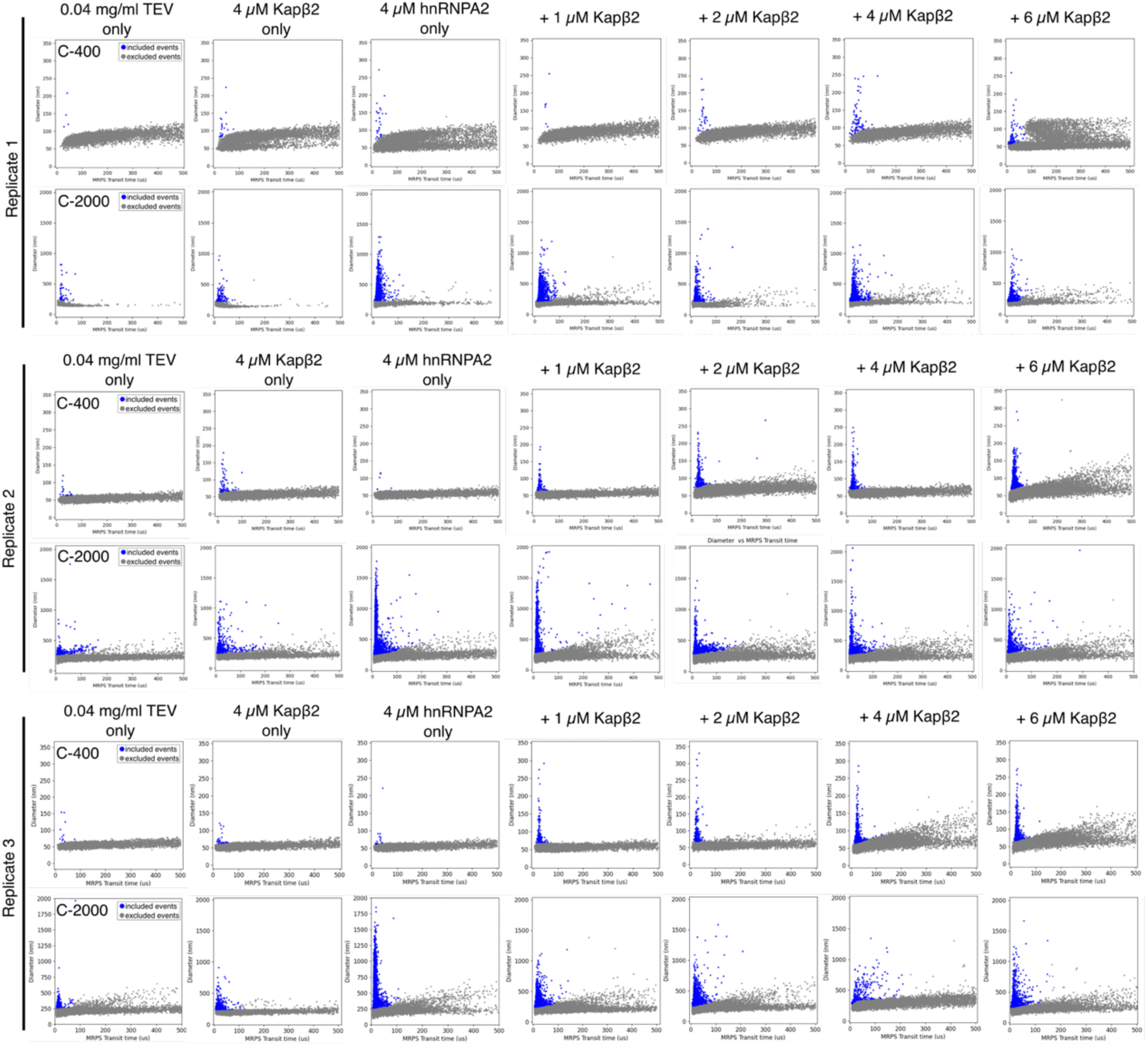
Raw MRPS scatter plots of hnRNPA2 in absence and presence of increasing Kapβ2 concentrations. Measurements were carried out as triplicates. Two different microfluidic cartridges (C-400 and C-2000) were used to cover a large range of particle sizes. 0.04 mg/ml TEV protease and 4 µM Kapβ2 alone were used as controls. Gray dots are the noise floor of the instrument. Blue dots represent single assemblies of distinct sizes which were used to extract size distributions depicted in Fig. 6d.

**Supplementary Fig. 12:**
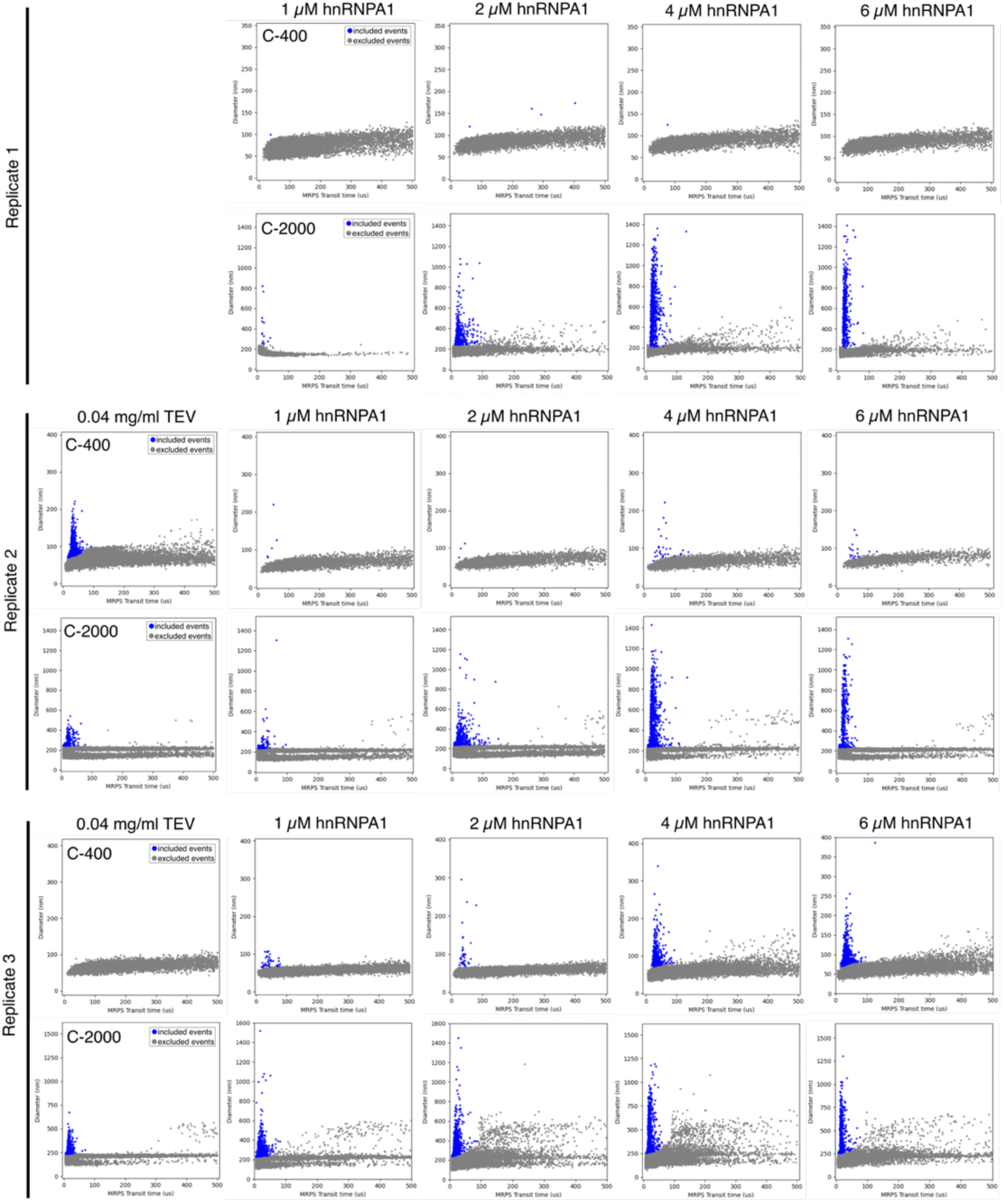
Raw MRPS scatter plots of samples at increasing hnRNPA1 concentrations. Plots depict the particle diameter in nm versus the MRPS transit time in s. Each sample was measured in three replicates. 0.04 mg/ml TEV protease served as control sample. Each blue dot represents a single event of a particle crossing the nano constriction in the respective cartridge (C-400 and C-2000). Size distributions corresponding to this data are shown in Extended Data Fig. 4b.

**Supplementary Fig. 13:**
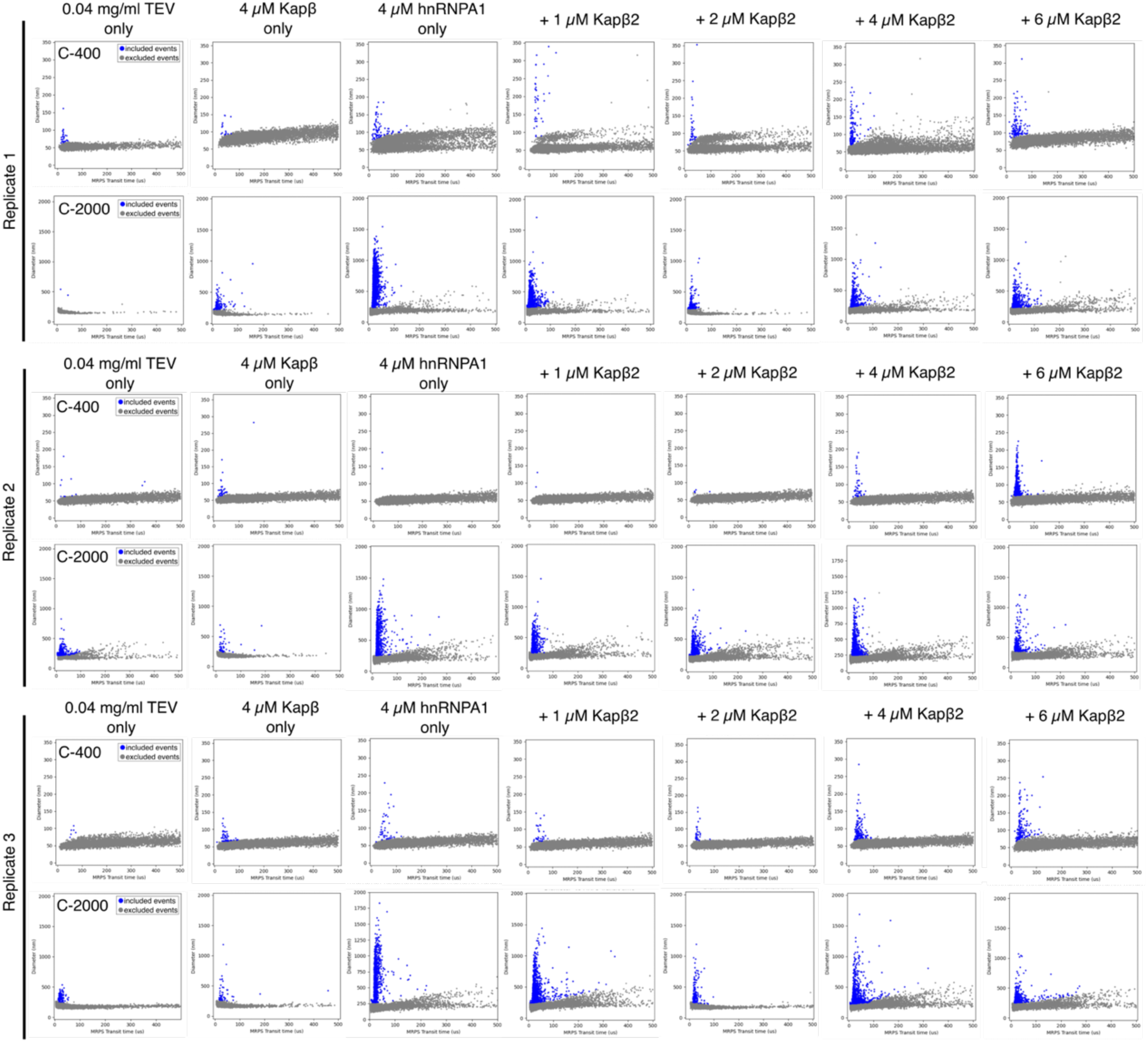
Raw MRPS scatter plots of hnRNPA1 in absence and presence of increasing Kapβ2 concentrations. Scatter plots show the particle diameter versus the MRPS transit time. Three independent replicates of each sample were measure. 0.04 mg/ml TEV protease and 4 µM Kapβ2 alone were analyzed as controls. All samples were measured using cartridges of two different sizes (C-400 and C-2000). Blue dots represent single assemblies, gray dots are the noise of the instrument and were excluded from the analysis. Corresponding size distributions are shown in Extended Data Fig. 4d.

**Supplementary Fig. 14:**
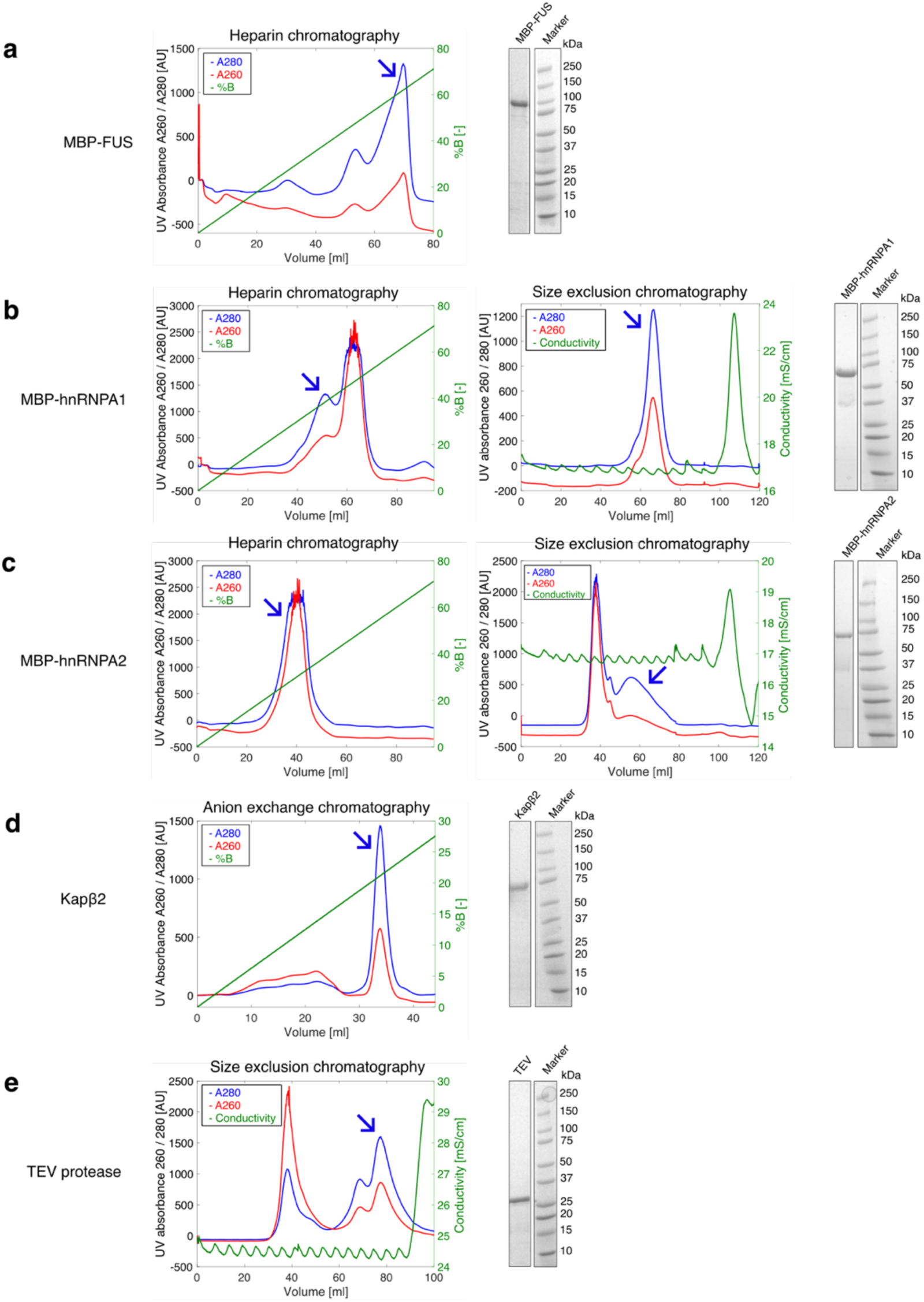
Purification of all proteins used in this study. a, Purification of MBP-FUS via Amylose-based affinity chromatography and Heparin chromatography (The right panel shows a representative SDS-PAGE demonstrating protein purity. **b,** MBP-hnRNPA1 was purified using amylose-based affinity chromatography, Heparin chromatography to remove bound RNA, and SEC. SDS-PAGE was used to confirm purity of final product. **c,** MBP-hnRNPA2 was purified using the same procedure as MBP-hnRNPA2. **d,** Kapβ2 was purified using Nickel-NTA affinity chromatography followed by anion exchange chromatography. SDS-PAGE confirms protein purity. **e,** TEV protease was purified via Ni-NTA affinity chromatography and SEC. The final protein product was pure, as shown via SDS-PAGE. In all panels, arrows indicate the peak containing the protein-of-interest for further purification or to be stored.

**Extended Data Fig. 1:**
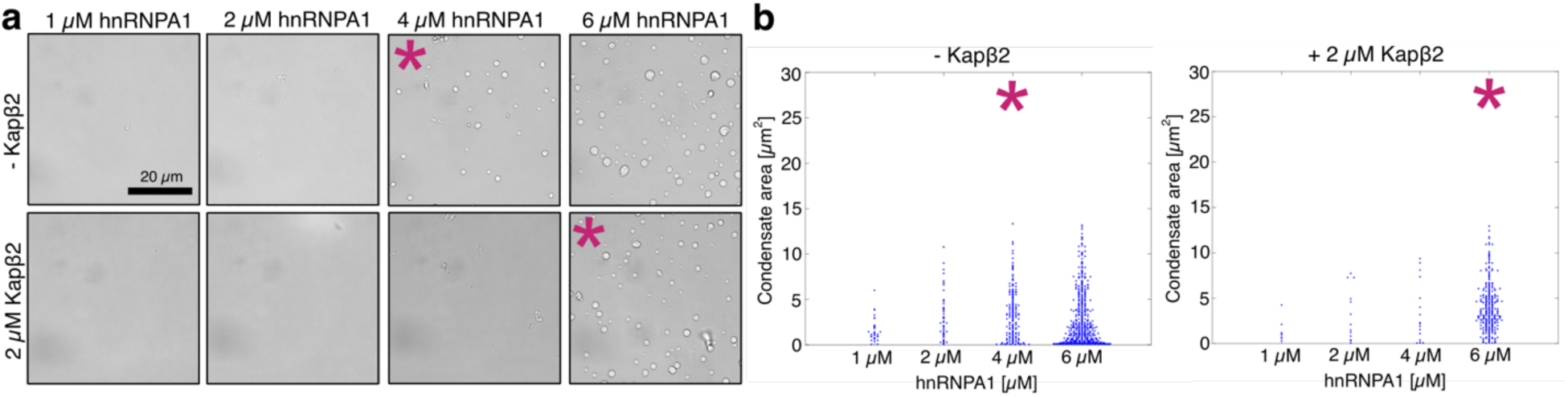
Kapβ2 shifts the saturation concentration *c*_sat_ of hnRNPA1 to higher values. **a,** Brightfield microscopy of samples at increasing hnRNPA1 concentrations in absence and presence of 2 µM Kapβ2. The intrinsic *c*_sat_ of 4 µM increases to 6 µM in the presence of Kapβ2. **b,** Quantification of the condensate area extracted from images in a. Asterisks (*) in a and b indicate *c*_sat_. Each blue dot corresponds to the area in µm^2^ of a single condensate, and the beeswarm plot is a combination of three images.

**Extended Data Fig. 2:**
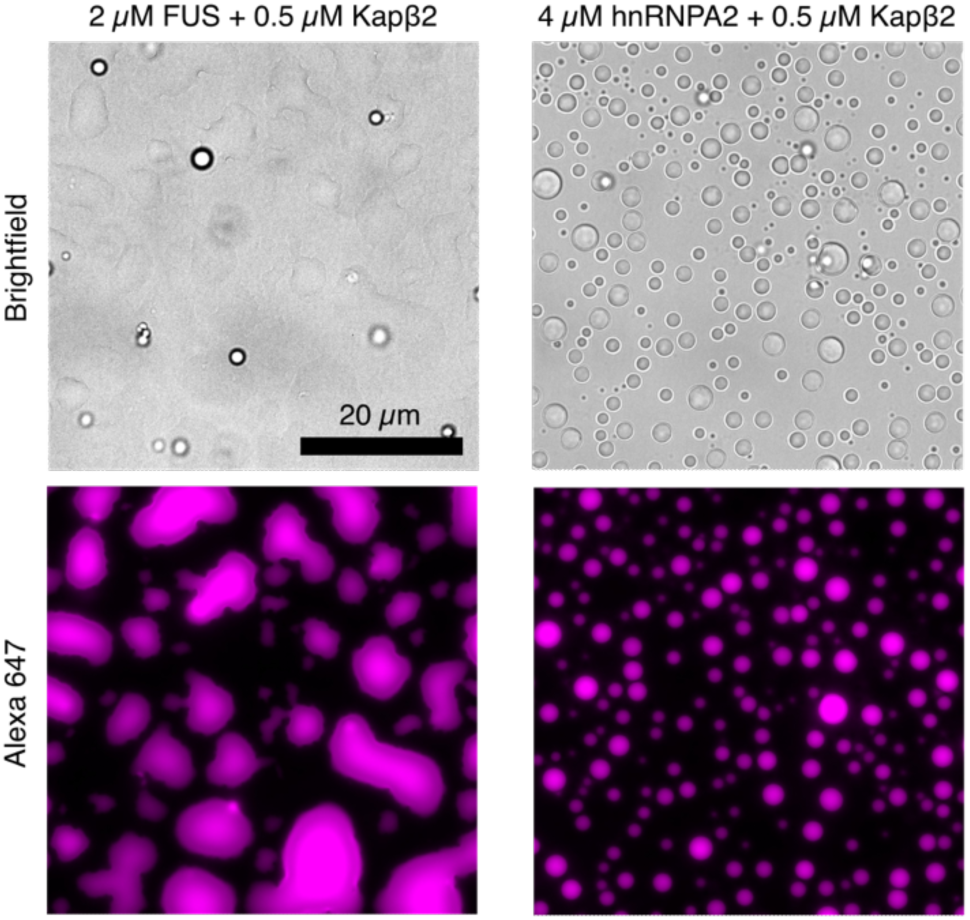
**Kap**β**2 partitioning into FUS and hnRNPA2 condensates.** 0.5 µM Alexa647-labeled Kapβ2 partitions into FUS (left panels) and hnRNPA2 condensates (right panels) formed at 2 and 4 µM protein, respectively.

**Extended Data Fig. 3:**
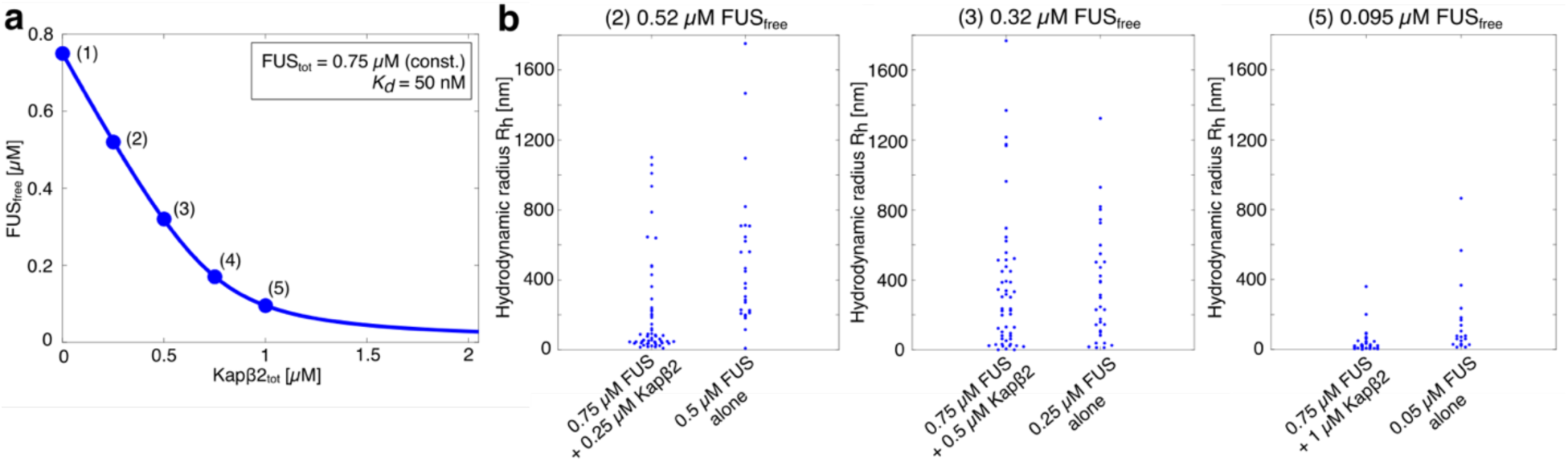
FUS_free_ is regulated by the binding of Kapβ2 and this in turn determines the spectrum of species that form via associations of FUS that are probed at low concentrations using FCS. a, Concentration of FUS_free_ as a function of the total Kapβ2 concentration (Kapβ2_tot_). Conditions (1)-(5) were experimentally tested via FCS. The concentration of FUS_tot_ was constant at 0.75 µM, a dissociation constant K_D_ of 50 nM was assumed for the Kapβ2:FUS complex. **b,** Comparison of hydrodynamic radii extracted using FCS of samples of FUS alone and samples containing 0.75 µM FUS with increasing concentrations of Kapβ2. Compared samples correspond to similar concentrations of FUS_free_.

**Extended Data Fig. 4:**
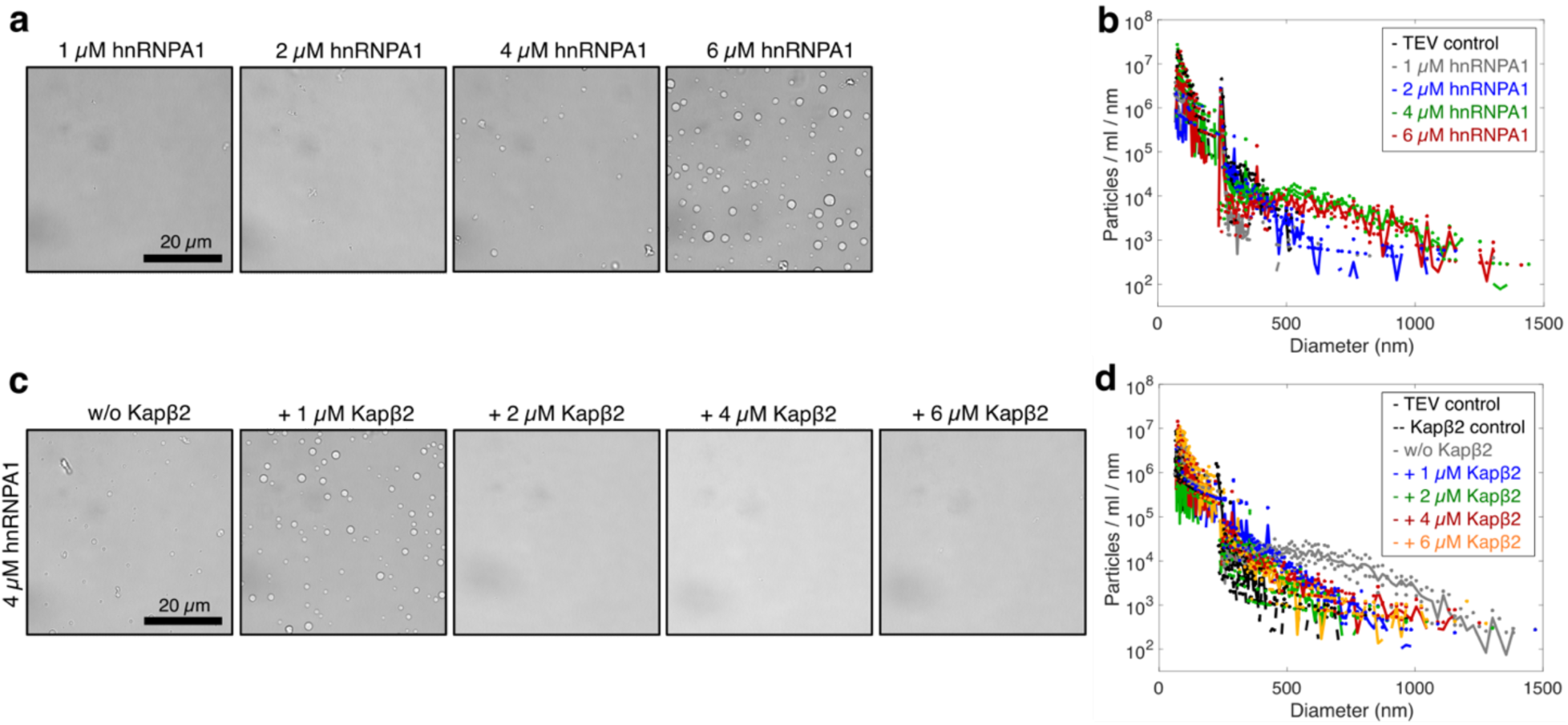
Kapβ2 inhibits hnRNPA1 self-assembly. **a**, Representative brightfield micrographs of samples at increasing concentrations of hnRNPA1. The saturation concentration *c*_sat_ is at 4 µM. **b,** Size distributions of samples at increasing hnRNPA1 concentrations show heavy-tailed character above *c*_sat_. **c,** Effect of Kapβ2 on hnRNPA1 phase separation. Sub-stoichiometric concentrations of Kapβ2 inhibit hnRNPA1 condensate formation. **d,** MRPS-measured size distributions of conditions depicted in c. Kapβ2 inhibits hnRNPA1 self-assembly. Size distributions lose their heavy-tailed character at concentrations at which Kapβ2 inhibits condensate formation (> 2 µM). Distributions in b and d show the mean particle counts/ml/nm of three independent replicates; dots represent the raw spread of the three replicates.

**Extended Data Fig. 5:**
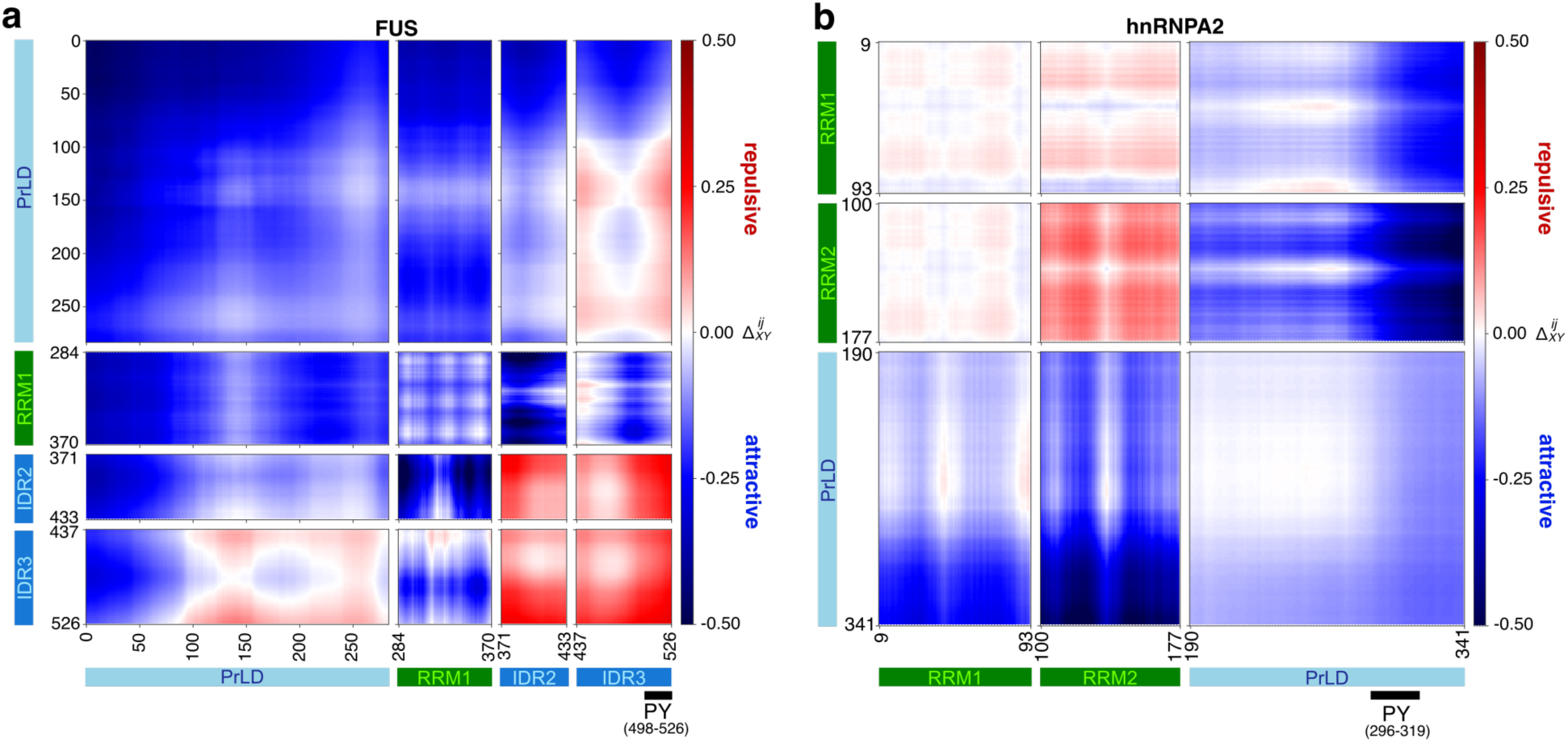
Comparison of the normalized inter-domain inter-residue distance maps of FUS and hnRNPA2 computed from atomistic simulations shown as Δ^ij^_XY_ reveal distinct interaction patterns. **a,** Inter-domain inter-residue distance map of FUS. The PrLD in FUS shows strong self-associations and preferential associations with the RRM and C-terminal R-rich regions, IDR2 and IDR3. In addition, the RRM in FUS also exhibits strong associations and with both IDR2 and IDR3. **b,** Inter-domain inter-residue distance map of hnRNPA2. hnRNPA2 shows weaker patterns of associations, mediated primarily by the PrLD where the inter-RRM interactions are repulsive or non-interacting. In both panels, the sequence architecture consisting of the simulated pair of domains X and Y for each protein are mapped to the axes. Regions of inter-residue attractions, Δ^CD^ < 0, are shown in blue whereas regions of inter-residue repulsions, Δ^CD^ > 0, are shown in red (see color scale bar).

